# A Cell-Autonomous Cold Sensor Promotes Lipolysis Through Phase Separation

**DOI:** 10.64898/2026.10.22.760470

**Authors:** Yong Chen, Li Chen, Zezhou Zhao, Tian Lu, Youchen Guan, Zinger Yang Loureiro, Shingo Kajimura

## Abstract

Temperature is a fundamental determinant of mammalian metabolism. The canonical cold-adaptation pathway involves cold sensing by peripheral sensory neurons, which activate sympathetic outflow, β-adrenergic signaling, and lipolysis in adipose tissue. In contrast, we report here a cell-intrinsic mechanism by which physiologically relevant cooling temperatures (27-31 °C) enhance lipolysis in subcutaneous adipose tissue. Mechanistically, cooling drives liquid-liquid phase separation (LLPS) of perilipin 1 (PLIN1), the dominant lipid droplet coat protein, promoting its condensation and the local recruitment of PKA to the droplet surface. This spatial reorganization selectively amplified PKA-mediated perilipin 1 phosphorylation, CGI-58 release, and ATGL-dependent lipolysis. The process depends on the intrinsically disordered region (IDR) of perilipin1, where four aromatic residues, conserved in endotherms but not in ectotherms, are required for cooling-induced LLPS and lipolysis in vivo. Notably, targeted delivery of an LLPS-competent perilipin 1 variant to the inguinal adipose tissue of mice enhanced local lipolysis and reduced depot mass. Together, these findings uncover a cell-intrinsic thermosensitive mechanism that can tune lipolytic responsiveness, offering a possible strategy for targeted depot-selective fat reduction.

## INTRODUCTION

Mammalian body temperature is thought to be maintained rigidly at an average of 37°C; however, it varies widely by anatomical location and dynamically fluctuates across days, seasons, and menstrual cycles (*1–4*). Notably, peripheral tissue temperatures can be lower than core temperature by 5°C or more (*4, 5*), and variations in wrist temperature amplitude are strong predictors of future disease risk, including type 2 diabetes, non-alcoholic fatty liver disease, chronic kidney disease, and hypertension (*6*). Despite this strong link between tissue temperature and metabolic health, the mechanisms by which physiologically relevant changes in peripheral tissue temperature directly influence metabolism remain poorly understood.

At the whole-body level, substantial progress has been made in elucidating adaptive thermogenic responses to cold temperatures, particularly to severe cold exposure (e.g., 4-8 °C) (*7, 8*). The best-characterized pathway involves cold detection by peripheral thermosensory neurons, transmission of the temperature changes to the central nervous system, and activation of sympathetic outflow (*9*). Norepinephrine (NE) released from sympathetic nerve terminals acts on β-adrenergic receptors, including the β3-adrenergic receptor (β3-AR) expressed in adipocytes, which triggers PKA-dependent signaling and lipolysis (*10, 11*). The liberated fatty acids derived from white adipose tissue (WAT) are key substrates for thermogenesis in brown adipose tissue (BAT) (*12–14*). At the molecular level, temperature sensing in afferent neurons is mediated by the transient receptor potential (TRP) channel family, which collectively spans the full perceptual temperature spectrum (*15, 16*). Among these, TRPM8 responds to cool temperatures (23-28 °C) and to menthol, with activity increasing as temperatures decline into the innocuous cool range (*17, 18*). When temperature falls to noxious levels (<17 °C), additional pathways, such as TRPA1-expressing nociceptors, are recruited in some species (*19, 20*). This sensory-sympathetic axis functions as an essential systemic adaptive pathway that triggers lipolysis in adipose tissue.

In contrast, far less is known about how physiologically relevant cooling temperatures within peripheral tissues directly influence cellular metabolism. A clue came from the following unexpected observations: we shaved the area over subcutaneous inguinal WAT in wild-type mice and applied a cold pad (< 6 °C) to one side and a warm pad (35-37 °C) to the contralateral side for 40 min (10 min x 4 cycles) per day at room temperature (**Figure 1A**). Infrared imaging confirmed a local decrease in tissue temperature on the cold-pad side relative to the warm control side (**Figure 1B**). After 5 days of treatment, the inguinal WAT mass on the cold-pad side was significantly lower than that on the control side within the same animals (**Figure 1C**). The effect of tissue cooling was consistent and reproducible in three independent cohorts, both in males (12-week-old and 26-week-old) and females (16-week-old) (**Supplementary Figure 1A, B**). We should note that this effect was not a handling artifact of the cold pads, because exposure to the warm pads on both sides of the subcutaneous region did not alter the inguinal WAT weight (**Supplementary Figure 1C**).

**Figure 1.**
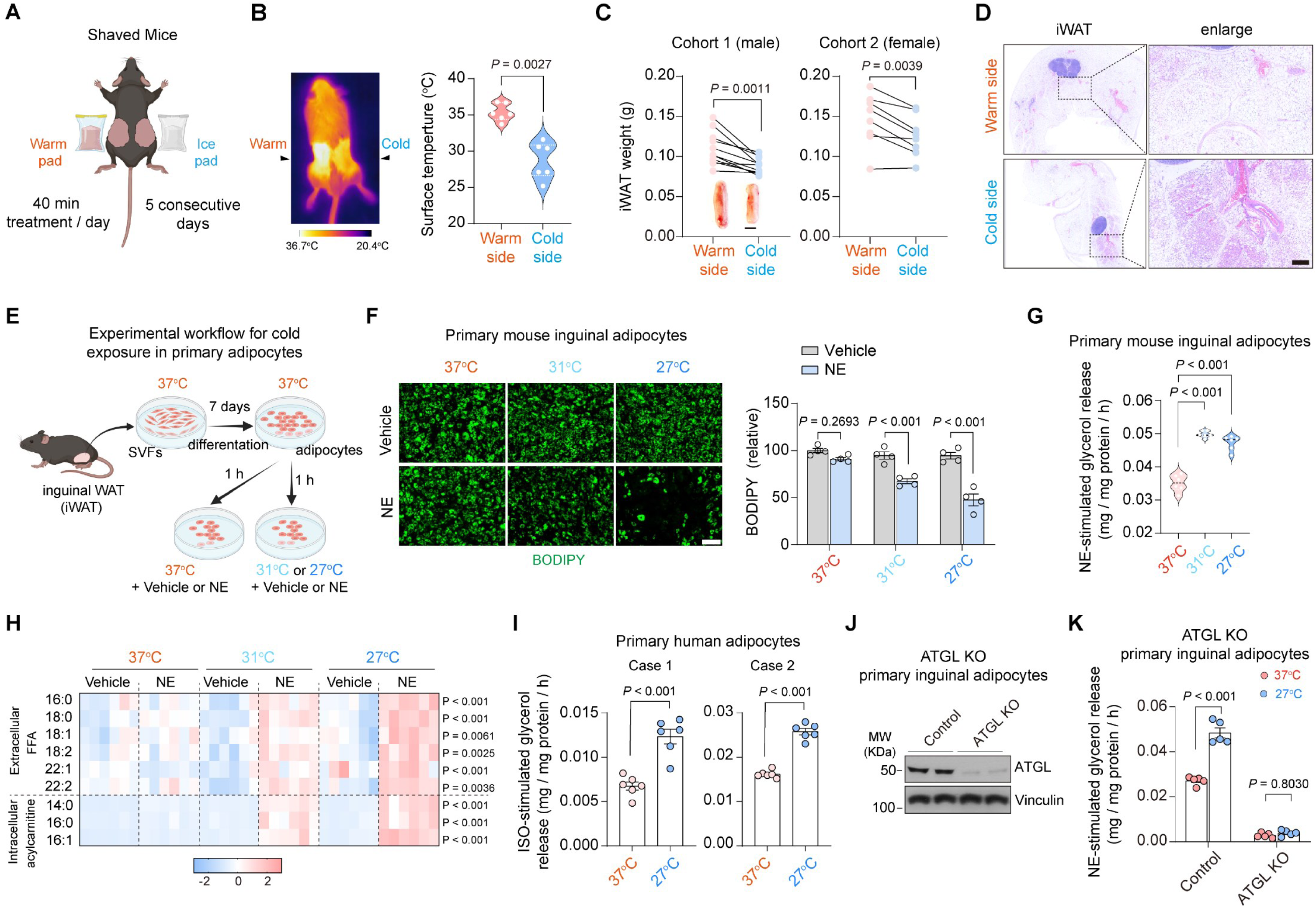
Cell-autonomous enhancement of lipolysis by cooling temperatures. **A.** Schematic illustration of the local cold exposure model in mice. Male or female mice housed at room temperature were shaved on both inguinal sides. Under anesthesia, mice were placed on either a warm or cold pad for 10 min, then allowed to recover on a warm pad in a lateral position until fully awake. This procedure was repeated 4 times per day for a total of 40 min of cold exposure per day for 5 consecutive days. **B.** <u>Left:</u> A representative whole-body thermal image obtained by infrared thermography after 3 min treatment with an ice pad or warm pad. <u>Right</u>: quantification of skin surface temperature from the left and right inguinal regions measured with an infrared thermometer. *n* = 6 mice. Data are mean ± s.e.m. Statistical significance: Paired *t*-test. **C.** Inguinal white adipose tissue (iWAT) weight from matched warm- and cold-treated depots within the same mouse. Each data point represents an individual depot, with paired measurements from the left and right sides connected. <u>Left</u>: 12-week-old male mice (*n* = 11). Representative iWAT images are shown. Scale bar: 0.5 cm. <u>Right</u>: 16-week-old female mice (*n* = 8). Statistical significance: Paired *t*-test. **D.** Representative hematoxylin and eosin (H&E) staining of iWAT sections from male mice in (**C**, left). <u>Left</u>, whole-slide scans; <u>Right</u>, magnified views of the indicated regions. Scale bar: 200 μm. **E.** Workflow of *in vitro* adipocyte cold exposure experiments. Stromal vascular fraction (SVF) cells from C57BL/6J mouse iWAT were differentiated for 7 days at 37 °C into mature adipocytes. Mature adipocytes were then incubated in a lipolysis medium (DMEM with 2% fatty acid-free BSA) at 37 °C, 31 °C, or 27 °C for 1 h, and subsequently stimulated with norepinephrine (NE, 0.1 μM) or vehicle for up to 10 h. **F.** <u>Left</u>: Representative BODIPY staining of mature adipocytes after 10 h NE treatment. Cells were stained with BODIPY for 10 min before imaging. <u>Right</u>: Quantification of BODIPY fluorescence intensity. *n* = 4 per group. Data are mean ± s.e.m. Two-way ANOVA with Šídák’s multiple comparisons test. **G.** Lipolysis assessed by glycerol release following NE stimulation for 10 h. Glycerol contents in the conditioned medium were normalized to the total protein of cells. *n* = 6 per group. One-way ANOVA followed by Tukey’s multiple comparisons test. **H.** Heatmap showing changes in BSA-bound free fatty acids (FFA) in the medium and cellular acyl-carnitines under the indicated conditions. Values are presented as Z-scores. *n* = 6 per group. *P* values indicate comparisons between the 27 °C + NE and 37 °C + NE groups. Unpaired *t*-test. **I.** Primary SVF cells isolated from subcutaneous adipose tissue of two human donors were differentiated into mature adipocytes and subjected to the cold-treatment protocol described in (**E**). Lipolysis was stimulated with isoproterenol (ISO, 5 μM) for 6 h. Glycerol levels in the conditioned media were quantified and expressed as ISO-stimulated glycerol release. *n* = 6 per group. Unpaired *t*-test. **J.** SVF cells isolated from the inguinal adipose tissue of *Pnpla2* flox mice were differentiated for 3 days and infected with adenovirus expressing iCre recombinase (Ad-iCre, ATGL KO) or GFP control (Ad-GFP, Control). Cells were differentiated for an additional 4 days, and ATGL protein levels were analyzed by western blotting. **K.** Mature adipocytes derived from (**J**) were subjected to the cold-treatment protocol described in (**E**). Lipolysis was assessed by NE-stimulated glycerol release over 6 h. *n* = 5 per group. Two-way ANOVA with Šídák’s multiple comparisons test.

Morphologically, the cooled WAT depot contained smaller adipocytes with multilocular lipid droplets and increased mitochondrial number, with elevated expression of brown/beige fat-selective genes relative to the control depot (**Figure 1D, Supplementary Figure 1D-G**). By contrast, tissue mass and morphology of interscapular BAT depots, which did not directly receive a cold or warm pad, were unchanged between the left and right lobes, consistent with preserved systemic body temperature under these conditions (**Supplementary Figure 1H-J**). In addition, we did not find notable differences in blood parameters, including fasting glucose, insulin levels, and ALT and AST levels, between the control group (warm pads on both sides) and the experimental group (a cold pad on one side and a warm pad on the other side) (**Supplementary Figure 1K-N**). Because the manipulation was spatially restricted to one side of the inguinal WAT within the same animal, these results suggested that local temperature can directly influence adipose remodeling independently of systemic factors, such as food intake or systemic neurohormonal and metabolic levels. We therefore set forth to determine whether subcutaneous adipose tissue possesses a cell-autonomous mechanism that enhances lipolysis in response to mild cooling.

## RESULTS

### Cell-autonomous enhancement of lipolysis by physiologically relevant cooling temperatures

To examine the extent to which the local cooling effect was mediated by adipocyte-intrinsic mechanisms, we isolated primary stromal vascular fractions (SVFs) from inguinal WAT of wild-type mice and differentiated them into adipocytes at 37 °C. Fully differentiated adipocytes were then equilibrated at 31 °C or 27 °C for 60 min, with 37 °C as a control, and subsequently stimulated with norepinephrine (NE) at 0.1 µM to induce lipolysis (**Figure 1E**). In the absence of NE, cooling did not alter intracellular triglyceride (TG) content as measured by BODIPY staining. Following NE stimulation, however, intracellular TG content decreased by 25.9% at 31 °C and by 47.7% at 27 °C relative to 37 °C (**Figure 1F**, quantification on the right).

The cooling-induced reduction in TG content suggested enhanced lipolysis in response to norepinephrine. In agreement, NE-induced glycerol release into the conditioned medium increased by 42% at 31 °C and by 35% at 27 °C (**Figure 1G**). Lipidomic analyses of the conditioned media further showed that cooling significantly increased the release of free fatty acids, including 16:0, 18:0, and 18:1 (**Figure 1H**). By contrast, intracellular acylcarnitine contents (14:0, 16:0, and 16:1) were significantly elevated at 27 °C following NE treatment, while the triglyceride/diacylglycerol (TG/DAG) ratio decreased (**Figure 1H, Supplementary Figure 2A**). These results indicate that NE-induced lipolysis is enhanced in adipocytes under cooled conditions. We also observed consistent results in primary and immortalized brown adipocytes, in which cooling to 27 °C enhanced NE-induced lipolysis (**Supplementary Figure 2B-F**). Importantly, human adipocytes derived from the subcutaneous WAT of two independent donors showed a greater isoproterenol-induced glycerol-release response at 27 °C than at 37 °C (**Figure 1I**). The results are in alignment with previous studies indicating that cooling affects anabolic or catabolic pathways in adipocytes (*21, 22*), although the underlying mechanism remains unknown.

To test the downstream pathway, we used two independent approaches to block adipose triglyceride lipase (ATGL; also known as PNPLA2), which is required for adipocyte lipolysis (*23, 24*). First, we genetically deleted ATGL in primary cultured adipocytes derived from inguinal WAT SVFs of *Pnpla2* flox mice by infecting differentiated adipocytes with Ad-iCre or Ad-GFP control (ATGL deletion in mature adipocytes) (**Figure 1J**). Cultured adipocytes were equilibrated at 37 °C or 27 °C for 60 min and then stimulated with NE. Cooling potentiated NE-induced lipolysis in control cells, as measured by glycerol release into the medium, whereas this effect was completely blunted in ATGL knockout (KO) adipocytes (**Figure 1K**). Second, pharmacological inhibition of ATGL with Atglistatin (*25*) similarly blunted the effect of cooling on NE-stimulated lipolysis (**Supplementary Figure 2G, H**). Together, these results show that physiologically relevant cooling temperatures enhance ATGL-dependent adipocyte lipolysis in a cell-autonomous manner.

### Cooling temperatures selectively amplify PKA signaling independent of the TRP channels

A general principle in biology is that lower temperatures slow many biochemical processes, also known as the Q10 effect (*26*). The observations above, however, suggested that adipocytes represent an exception in which cooling enhances NE-induced lipolysis. At the transcriptional level, cooling with NE stimulation did not substantially upregulate the expression of β-ARs (*Adrb1, Adrb2, Adrb3*), PKA (*Prkaca*, *Prkar2b*), *Plin1*, or ATGL (*Pnpla2*) (**Supplementary Figure 3A**). We then used phosphoproteomics to map how cooling alters NE-induced signaling (**Figure 2A**). The analysis found that 4 phosphopeptides were significantly increased at 27 °C in response to NE, whereas 19 phosphopeptides were decreased (**Figure 2B, Supplementary Figure 3B, Table S1**). Among them, phosphorylation of perilipin 1 at S81 and S492 caught our attention because perilipin 1 is the dominant lipid-droplet protein in adipocytes and a key regulator of lipolysis through hormone-stimulated activation of ATGL and hormone-sensitive lipase (HSL) (*27–31*). Notably, all of the cooling-enhanced phosphorylation sites, including perilipin 1^S492^, perilipin 1^S81^, TNKS1BP1^S429^, and CEP170^S950^, contained consensus motifs for protein kinase A (PKA) (**Figure 2C**). This aligns with the established role of PKA in phosphorylating perilipin 1 at six sites, including the C-terminal site S492, which is important for adipose lipolysis (*32, 33*). We also detected phosphorylated HSL (S34, S559, S597, S651, and T752), another well-known PKA target in adipocytes, as well as additional phosphorylated sites in perilipin 1 (S384 and S460), although these were similar between 27 °C and 37 °C (**Table S1**). These results suggest that cooling does not stimulate all downstream PKA targets; rather, it amplifies a subset of them. In agreement, immunoblotting with a PKA-substrate antibody showed that cooling to 27 °C selectively increased the PKA-phosphorylated signal at approximately 55kDa (indicated by an arrowhead) that corresponded to perilipin 1, within 5 min after NE treatment (**Figure 2D**). This observation was consistent across three other models of differentiated adipocytes, including primary inguinal WAT-derived adipocytes, immortalized inguinal WAT-derived adipocytes, and immortalized brown adipocytes (**Supplementary Figure 3C, D, E**).

**Figure 2.**
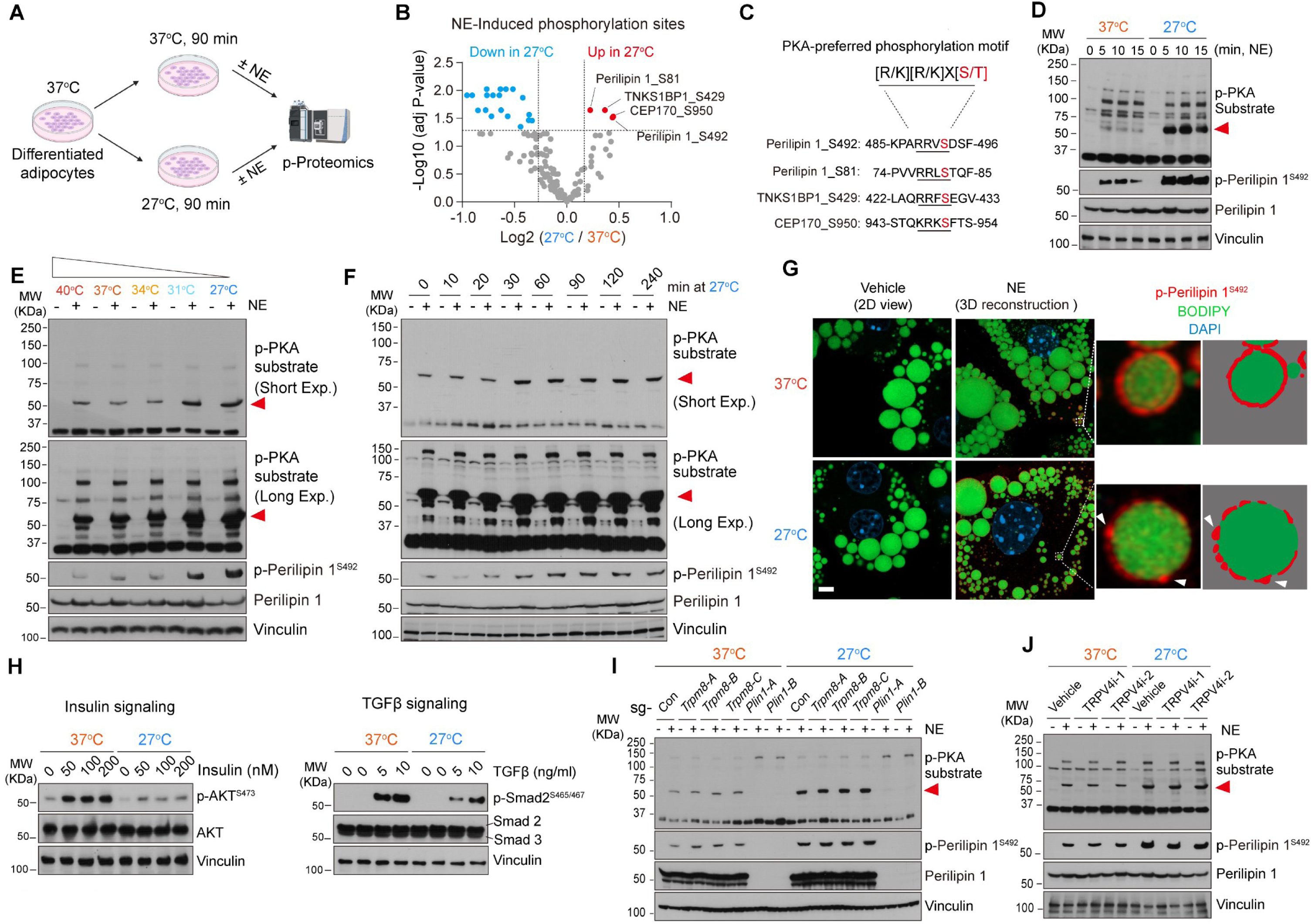
Cooling temperatures selectively amplified PKA signaling independent of the TRP channels. **A.** Schematic overview of the phosphoproteomics workflow. Immortalized brown preadipocytes were differentiated for 6 days at 37 °C and incubated at either 37 °C or 27 °C for 90 min. Cells were subsequently stimulated with norepinephrine (NE) at 0.1 μM or vehicle (PBS) for 10 min, rapidly harvested, and processed for phosphoproteomic analysis. **B.** Volcano plot showing NE-induced phosphorylation sites at 37 °C vs. 27 °C. Each dot represents an individual phosphorylation site. Red dots indicate sites that significantly increased at 27 °C, whereas blue dots indicate sites that decreased at 27 °C. The x-axis represents fold change (27 °C + NE / 37 °C + NE), and the y-axis indicates adjusted P values. **C.** Sequence motif analysis of cold-enhanced phosphorylation sites in (**B**). The enriched motif indicates consensus with the canonical PKA phosphorylation motif. **D.** Immunoblot analysis of time-dependent NE-induced phosphorylation. Differentiated adipocytes cultured at 37 °C or 27 °C were stimulated with NE at 0.1 μM for the indicated time points (5, 10, and 15 min), followed by immunoblot detection of phospho-PKA substrates, phospho-perilipin 1^S492^, and total perilipin 1. Vinculin was used as a loading control. The red arrowhead indicates p-Perilipin 1 at approximately 55 kDa. **E.** Temperature-dependent activation of PKA signaling. Differentiated brown adipocytes were incubated at 40 °C, 37 °C, 34 °C, 31 °C, or 27 °C for 90 min prior to NE stimulation at 0.1 μM for 10 min. Phosphorylation levels of PKA substrates, phospho-perilipin 1^S492^, and total perilipin 1 were analyzed by immunoblotting. The red arrowhead indicates p-Perilipin 1 at approximately 55kDa. **F.** Time-dependent effect of cold exposure on PKA signaling. Differentiated adipocytes were incubated at 27 °C for the indicated durations and stimulated with NE at 0.1 μM for 10 min. Phosphorylation of PKA substrates, phospho-perilipin 1^S492^, and total perilipin 1 were analyzed by immunoblotting. Vinculin was used as a loading control. **G.** Immunofluorescence analysis of phospho-perilipin 1^S492^. Differentiated primary inguinal white adipocytes were incubated at 27 °C for 90 min, followed by NE stimulation for 10 min. Cells were stained for phospho-perilipin 1^S492^ (Red), BODIPY (lipid droplet, Green), and DAPI (nuclei, Blue). Arrowheads show condensates of phospho-perilipin 1^S492^ signals. The right illustration was traced based on the 3D-reconstructed images. Scale bar: 5 µm. **H.** Temperature-dependent signaling in response to insulin and TGFβ. Differentiated brown adipocytes were cultured at 27 °C for 90 min and treated with increasing concentrations of insulin (left) or TGF-β (right) for 10 min. Activation of insulin signaling and TGF-β signaling was assessed by immunoblot detection of phospho-AKT^473^ and phospho-SMAD2^S465/467^, respectively. **I.** The requirement of TRPM8 and perilipin 1 for cold-enhanced PKA signaling. Brown preadipocytes transduced with lentiviral sgRNAs targeting TRPM8, perilipin 1, or sgControl were differentiated at 37 °C and incubated at 27 °C for 90 min, and then stimulated with NE at 0.1 μM for 10 min. The indicated phospho-proteins were examined by immunoblotting. Vinculin was used as a loading control. **J.** The requirement of TRPV4 for cold-induced PKA signaling. Differentiated brown adipocytes were treated with two independent TRPV4 inhibitors (HC-067047, GSK205) at either 37 °C or 27 °C for 90 min, followed by NE stimulation for 10 min. The indicated phospho-proteins were examined by immunoblotting. Vinculin was used as a loading control.

To define the adipocyte-intrinsic response, we next incubated differentiated adipocytes at 40 °C, 37 °C, 34 °C, 31 °C, or 27 °C for 90 min. We found that cooling to 31 °C or lower enhanced NE-induced phosphorylation of perilipin 1^S492^ (**Figure 2E**). This temperature threshold was well within the physiological temperature range observed in subcutaneous WAT (see Figure 1B). Time-course experiments further showed that exposure to 27 °C for 30 min or longer was sufficient to enhance NE-induced phosphorylation of perilipin 1^S492^ (**Figure 2F**). In turn, the cooling-enhanced p-perilipin 1 was attenuated to a basal level when rewarmed to 37°C for 60 min or longer (**Supplementary Figure 3F**). The following observations suggest that the effect of cooling on p-perilipin 1^S492^ reflected enhanced PKA signaling, rather than altered β-AR abundance or cAMP turnover: First, cooling itself had no effect on intracellular cAMP levels (**Supplementary Figure 3G**), whereas it enhanced perilipin 1^S492^ phosphorylation in the presence of a cAMP analog 8-Br-cAMP (**Supplementary Figure 3H**). Second, the PKA inhibitor H89 completely blocked the cooling-induced increase of p-perilipin 1^S492^, whereas the phosphatase inhibitor okadaic acid did not blunt the action (**Supplementary Figure 3I, 3J**). Next, we performed immunostaining for p-perilipin 1^S492^ by confocal microscopy and, as expected, we observed a stronger signal in NE-treated adipocytes. Notably, three-dimensional (3D)-reconstructed image analysis revealed that p-perilipin 1 signals were distributed relatively uniformly around lipid droplets at 37 °C; however, at 27 °C, they frequently appeared as puncta-like structures on lipid droplets (arrowheads in the inset, **Figure 2G**). This observation is consistent with the pioneering work (*34, 35*) and also provides an important clue to the underlying mechanism. We will discuss this later in Figure 3.

**Figure 3.**
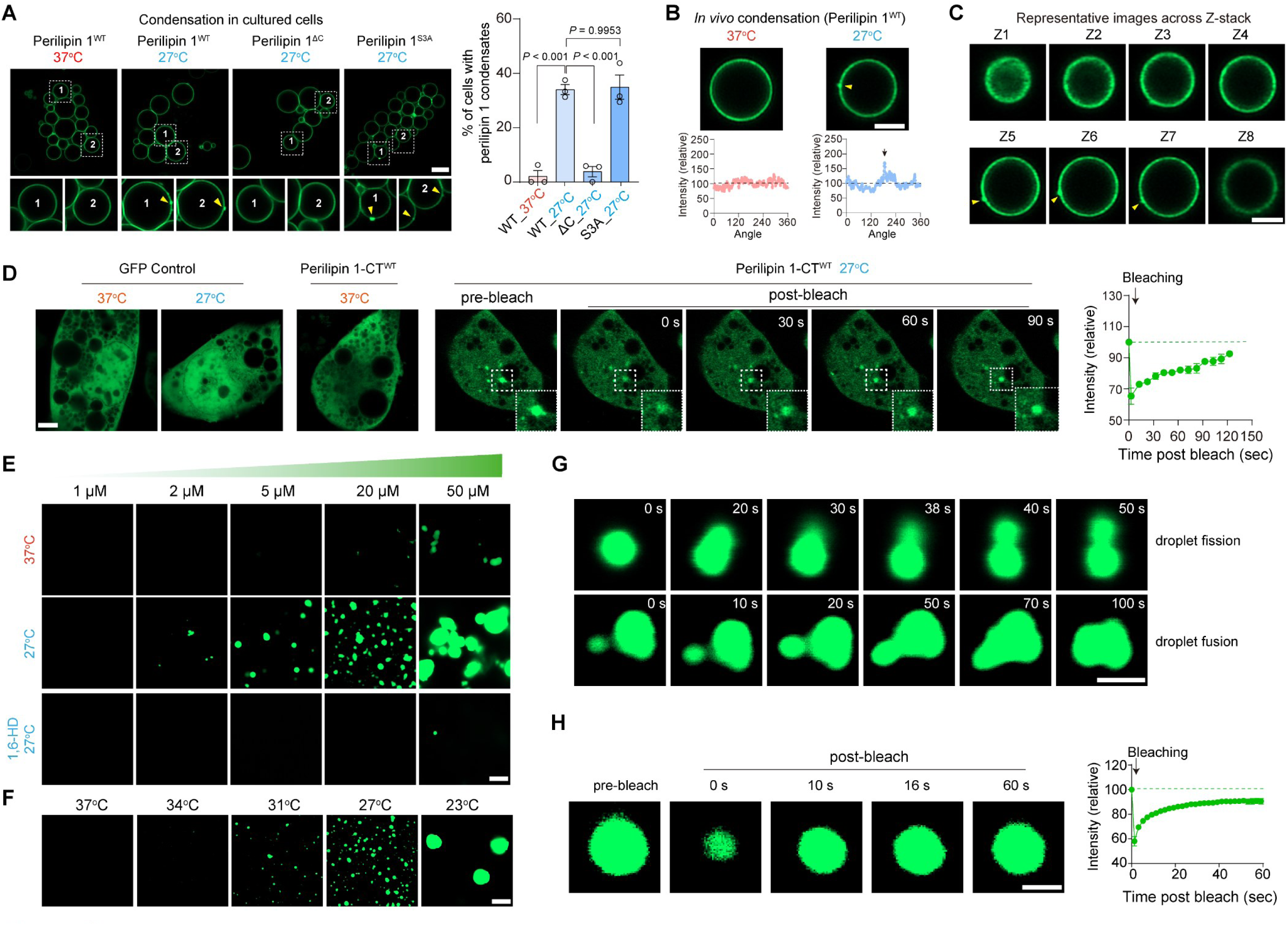
Cooling temperatures induce perilipin 1 phase separation. **A.** Cooling-induced condensate formation of perilipin 1 on lipid droplets. <u>Left</u>: Live-cell imaging analysis of GFP-perilipin 1^WT^, GFP-perilipin 1^ΔC^, and GFP-perilipin 1^S3A^ (C-terminal PKA phosphorylation defective mutant, S433/492/517A) in differentiated adipocytes at 37 °C or 27 °C. Insets show two magnified views of the boxed regions. Yellow arrows indicate punctate GFP signals formed on lipid droplets. Scale bar: 4 µm. <u>Right</u>: Percentage of cells containing perilipin 1 condensates. *n*= 3 independent experiments; each dot represents ≥ 15 cells. One-way ANOVA followed by Tukey’s multiple comparisons test. **B.** Cooling-induced condensate formation of perilipin 1 *in vivo*. <u>Upper panel</u>: Representative GFP-perilipin 1 images of isolated lipid droplets from iWAT incubated at 37 °C (left) or 27 °C (right). Arrowheads indicate perilipin 1 condensates on lipid droplet surfaces. Scale bar: 2 μm. <u>Lower panel</u>: Corresponding angular profiles of GFP fluorescence intensity on lipid droplet surfaces, quantified using Fiji software. **C.** Representative Z-stack images of the lipid droplet shown in (**B**). Arrowheads indicate perilipin 1 condensates on lipid droplets. Scale bar: 2 μm. **D.** The C-terminus of perilipin 1 is sufficient for condensate formation in adipocytes. Differentiated brown adipocytes stably expressing GFP fused to the C-terminus of perilipin 1 (perilipin 1-CT) or GFP alone as a control were incubated at 27 °C or 37 °C for 8 h. Representative fluorescence images show signal recovery before and after photobleaching (FRAP). Scale bar: 4 µm. <u>Right</u>, fluorescence recovery curves quantifying signal recovery within the bleached region after photobleaching. *n* = 5. **E.** *In vitro* phase separation assay with purified perilipin 1 C-terminal protein fused with GFP (GFP-perilipin 1-CT). GFP-perilipin 1-CT protein at the indicated concentrations was dissolved in buffer containing 150 mM NaCl, 25 mM HEPES (PH 7.4), and 1 mM DTT, and incubated with 7.5% PEG-8000 for 10 min at either 37 °C or 27 °C. 1,6-HD treatment at 10% was added together with PEG-8000. Scale bar: 4 µm. **F.** Temperature-dependent phase separation with purified GFP-perilipin 1-CT protein. Purified GFP-perilipin 1-CT protein at the concentration of 10 μM was prepared as (**E**) and incubated at 37 °C, 34 °C, 31 °C, 27 °C, and 23 °C. Scale bar: 4 µm. **G.** Perilipin 1 condensates exhibit dynamic fusion and fission behavior. Time-lapse imaging showing fission (top) and fusion (bottom) of GFP-perilipin 1-CT droplets at 23 °C. Scale bar: 1 µm. **H.** FRAP analysis of GFP-perilipin 1-CT droplets at 27 °C. Representative photobleaching images of GFP–perilipin 1-CT droplets were captured at the indicated time. Scale bar: 0.5 µm. Right, fluorescence recovery curves quantifying signal recovery of perilipin 1 condensates after photobleaching. *n* = 4.

These findings raised the question of whether the cooling effect is selective across hormone-induced signaling pathways. We therefore examined how cooling altered responses to insulin and TGFβ. In differentiated adipocytes, insulin-induced P-AKT^S473^, a canonical insulin-signaling readout, was markedly reduced at 27 °C relative to 37 °C (left panels in **Figure 2H**). Similarly, TGFβ-induced Smad2 phosphorylation at S465/467 was reduced at 27 °C compared with 37 °C (right panels in **Figure 2H**). Thus, cooling does not broadly enhance signaling in adipocytes; instead, it preferentially enhances the adrenergic PKA-perilipin 1 axis while attenuating insulin- and TGFβ-dependent signaling.

The best-known cold sensor is the TRP channel family, especially TRPM8 (*17, 18*). TRPM8 is expressed in brown and white adipocytes, and activation of TRPM8 by menthol has been reported to stimulate UCP1 expression (*36, 37*). Thus, we tested whether TRPM8 is required for cell-autonomous cooling-enhanced PKA signaling by generating CRISPR-based TRPM8 KO adipocytes using three independent gRNAs (**Supplementary Figure 4A**). We also generated CRISPR-*Plin1* KO cells using two independent gRNAs to confirm that the 55 kDa signal was indeed perilipin 1. Cooling to 27 °C robustly enhanced PKA-mediated perilipin 1^S492^ phosphorylation in TRPM8 KO adipocytes after NE treatment, to a level comparable to that in control adipocytes (**Figure 2I**). Consistently, cooling augmented NE-induced lipolysis in TRPM8 KO adipocytes, as measured by glycerol release and reduced intracellular lipid content (**Supplementary Figure 4B, C**). Similarly, pharmacological inhibition of TRPM8 did not affect cooling-enhanced PKA signaling (**Supplementary Figure 4D**). As an additional candidate, we examined TRPV4 using two specific inhibitors (GSK205 and HC-067047), as previous studies showed that cooling directly activated the expression of thermogenic genes, including *Ucp1*, in cultured adipocytes and that TRPV4 negatively regulates these thermogenic genes (*21, 38*). However, pharmacological inhibition of TRPV4 did not affect the cooling effect on NE-induced perilipin 1^S492^ phosphorylation (**Figure 2J**). These results suggest that cooling temperatures selectively amplify NE-induced PKA signaling to phosphorylate perilipin 1 and lipolysis, independent of TRPM8 and TRPV4.

### Cooling induces phase separation of perilipin 1 protein

Perilipin 1 is one of the most abundant lipid droplet-coating proteins in adipocytes and plays a central role in lipolysis by controlling lipase access to stored triglycerides and in regulating lipid storage (*27–31, 39*). The observation that the puncta-like p-perilipin 1^S492^ signal emerged on lipid droplets at 27 °C (Figure 2G) suggested that cooling might induce perilipin 1 protein condensation. Of note, *in silico* structural analyses using two independent algorithms, PONDR VSL2 and VLXT (*40*), predicted intrinsically disordered regions (IDRs) within mouse perilipin 1, including its C-terminal region (aa 406-517) (**Supplementary Figure 5A**). Human perilipin 1 also contains a C-terminal IDR, in agreement with a recent report (*41*). AlphaFold 3 predicted this region to be highly unstructured, despite differences in shape between mice and humans (**Supplementary Figure 5B**). Given the importance of IDRs in liquid-liquid phase separation (LLPS) across diverse biological contexts, including temperature (*42–48*), we asked whether perilipin 1 undergoes temperature-dependent phase separation. To test this, we first generated constructs encoding full-length perilipin 1 (aa 1-517) fused to GFP at the N-terminus, as well as a GFP-perilipin 1 mutant lacking the C-terminal domain (aa 1-405) (**Supplementary Figure 5C**). Subsequently, we expressed them in adipocytes and examined their localization on lipid droplets using live-cell confocal microscopy. We found that full-length perilipin 1 (perilipin 1^WT^) was localized to the surface of lipid droplets and was distributed relatively evenly at 37 °C regardless of NE treatment (**Figure 3A** – left panel, **Supplementary Figure 5D**). By contrast, incubation at 27 °C altered the distribution of perilipin 1, producing numerous puncta-like condensates on lipid droplets (indicated by arrows in **Figure 3A**). The 3D image reconstruction validated the presence of these condensates on lipid droplets (**Supplementary Figure 5E**). This cooling-induced condensate formation required the C-terminal region of perilipin 1, as the C-terminal deletion mutant (perilipin 1^ΔC^) did not form puncta at 27 °C. On the other hand, a phosphorylation-deficient mutant at the C-terminus (S433/492/517A; perilipin 1^3SA^) was still able to form condensates at 27 °C (**Figure 3A** – left panel, **Supplementary Figure 5F**), suggesting that cooling-induced perilipin 1 condensation does not require its phosphorylation. Quantification of the imaging data is shown in the right panel of **Figure 3A**. We also found that treatment with 1,6-hexanediol (1,6-HD), which disrupts weak hydrophobic interactions often required for LLPS, dispersed the cooling-induced perilipin 1 condensates, and these condensates rapidly re-formed after 1,6-HD withdrawal (**Supplementary Figure 5G**). Live imaging showed that perilipin 1 condensates emerged after 60 min or longer incubation at 27 °C (**Supplementary Figure 5H**), which aligns with the time-course change in the cooling-induced amplification of perilipin 1 phosphorylation.

To further determine whether cooling-induced perilipin 1 condensation occurs *in vivo*, we developed a system in which full-length GFP-perilipin 1 was expressed via AAV in the inguinal WAT of 8-week-old wild-type mice. Three weeks after injection, lipid droplets were isolated from the inguinal WAT and plated on a glass-bottom dish at 37 °C or 27 °C. Perilipin 1 localization on lipid droplets was then immediately visualized by live confocal imaging, together with angular distribution analysis, as validated in adipocytes (**Supplementary Figure 6**). Consistent with the results in cultured adipocytes, we observed robust punctate perilipin 1 signals on lipid droplet surfaces at 27 °C, but not at 37 °C (**Figure 3B**; top, representative GFP-perilipin 1 images; bottom, angular distribution of GFP fluorescence intensity). To validate these condensates, we acquired Z-stack images of perilipin 1 on lipid droplets (**Figure 3C**).

Next, we asked whether the C-terminal region of perilipin 1 is sufficient to form temperature-dependent condensates. To test this, we expressed the C-terminal region of perilipin 1 (aa 406-517; perilipin 1-CT), fused to GFP at the N-terminus, or GFP alone in adipocytes and transferred the cells to 27 °C or 37 °C. Cooling did not alter the distribution of GFP alone. By contrast, perilipin 1-CT was diffusely distributed in the cytosol at 37 °C but formed condensates at 27 °C (**Figure 3D**). Fluorescence recovery after photobleaching (FRAP) showed recovery of the perilipin 1-CT signal at 27 °C, indicating that these condensates remained mobile and dynamically reorganized (**Figure 3D**). In addition, treatment with 1,6-HD reversibly blocked the formation of perilipin 1-CT condensates (**Supplementary Figure 7A**). Of note, the C-terminal domain of perilipin 1 was sufficient to drive condensate formation at 27°C in non-adipocytes (HEK293 and AML12 cells), and these condensates were disrupted by 1,6-HD treatment (**Supplementary Figure 7B, C**). The FRAP experiments further demonstrated that these condensates were mobile and dynamically reorganized (**Supplementary Figure 7D**).

To test this in a cell-free reconstituted system, we next purified the C-terminal domain of perilipin 1 (**Supplementary Figure 7E**) and incubated the protein with 7.5% PEG-8000, following an established *in vitro* phase-separation assay (*49*). Cooling to 27 °C for 10 min promoted perilipin 1 condensation in a dose-dependent manner, and this was potently blocked by 1,6-HD (**Figure 3E**). Consistent with the temperature dependence of cooling-enhanced lipolysis, low temperatures at 31 °C or lower induced perilipin 1 condensation (**Figure 3F**) – well within the physiological temperature range observed in subcutaneous WAT. High-resolution imaging captured fusion and fission of perilipin 1 droplets over time, supporting liquid-like behavior (**Figure 3G**). Furthermore, FRAP experiments showed rapid recovery of fluorescent signal after photobleaching, suggesting that perilipin 1 condensates are dynamic (**Figure 3H**). Together, these results show that physiologically relevant cooling (27-31 °C) induces liquid-liquid phase separation (LLPS) of perilipin 1 through its C-terminal IDR.

### The structural requirement for cooling-induced LLPS of perilipin 1

From an evolutionary perspective, the C-terminal IDR of perilipin 1 is highly conserved among endotherms and contains several aromatic residues (F and Y). By contrast, ectotherms, including reptiles and fish, lack these domains (**Figure 4A**). The C-terminal IDR is unique to PLIN1 as it is absent in PLIN2, PLIN3, PLIN4, and PLIN5 (**Supplementary Figure 8A**). It is worth clarifying that perilipin 1 is by far the most dominant perilipin among perilipins expressed in mature adipocytes of both inguinal WAT and BAT (*50*), in alignment with publicly available transcriptional data (*51, 52*) (**Supplementary Figure 8B**). Furthermore, cooling did not induce condensate formation of perilipin 2, 3, or 5, nor did it alter the expression levels of any of these proteins in the inguinal WAT (**Supplementary Figure 8C**, **D**). Since previous studies reported increased perilipin 5 expression in BAT by systemic cold exposure (*53, 54*), the effect of local WAT cooling is distinct from that of systemic cold exposure in regulating perilipin 5.

**Figure 4.**
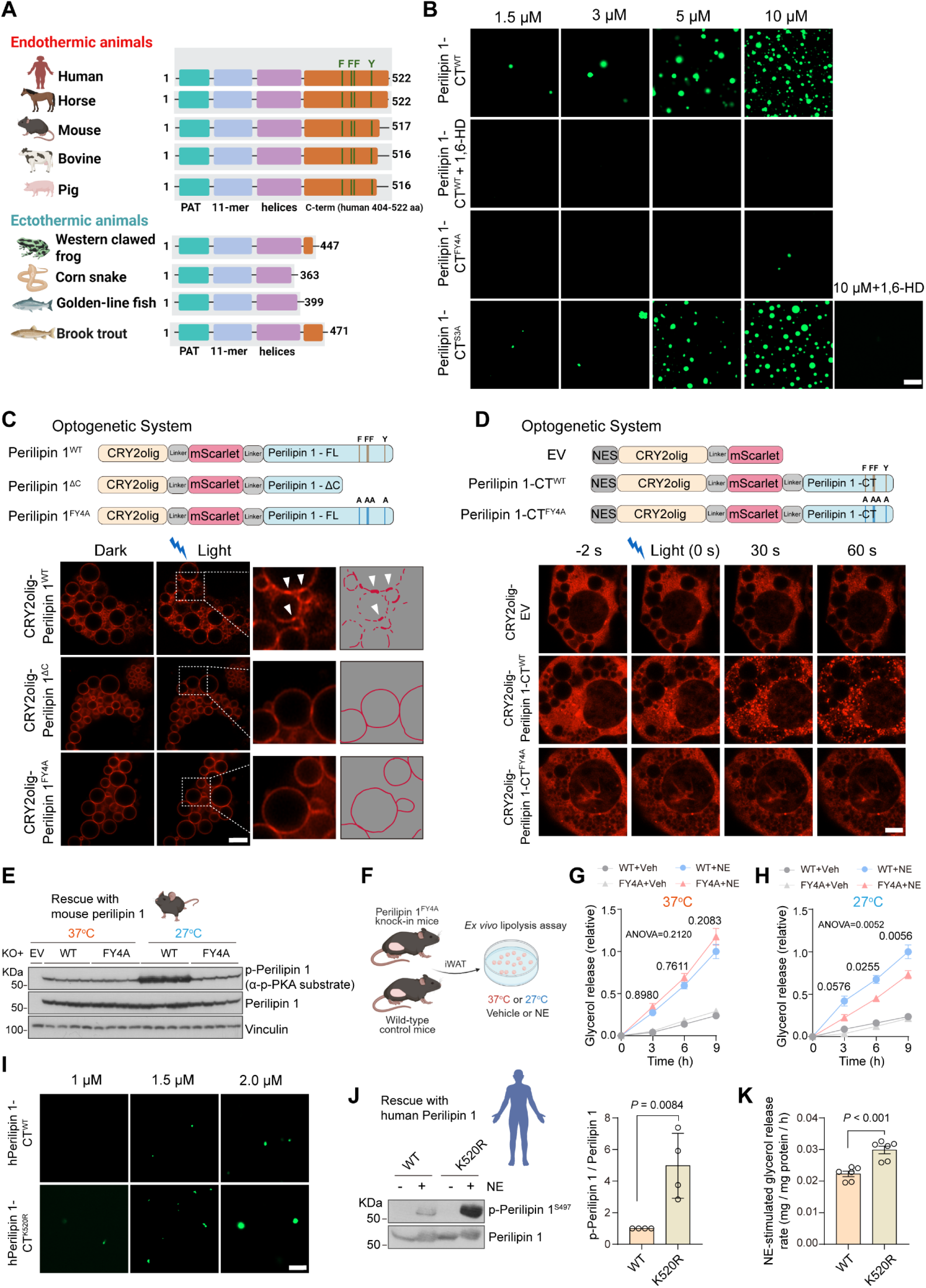
The structural requirement for cooling-induced perilipin 1 phase separation. **A.** Schematic comparison of perilipin 1 protein architecture across representative endothermic (human, horse, mouse, bovine, and pig) and ectothermic (western clawed frog, corn snake, golden-line fish, and brook trout) species. Conserved domains, including the PAT domain, 11-mer repeats, and amphipathic helices, are present in all species, whereas a C-terminal extension is selectively observed in endothermic animals. The C-terminal region contains conserved phenylalanine (F) and tyrosine (Y) residues, which are absent in ectothermic species. Amino acid lengths are indicated on the right. **B.** *In vitro* phase separation assays using purified GFP-tagged perilipin 1 C-terminal proteins (GFP-perilipin 1-CT, perilipin 1-CT^FY4A^, and perilipin 1-CT^S3A^) at the indicated concentrations in the absence or presence of 10% 1,6-HD. Scale bar: 4 µm. **C.** <u>Upper panel</u>: Schematic of the CRY2-based optogenetic system using the full-length perilipin 1 (perilipin 1^WT^), C-terminal truncation (perilipin 1^ΔC^), and phase separation-deficient mutant (perilipin 1^FY4A^). <u>Lower panels</u>: Differentiated brown adipocytes expressing the indicated constructs were imaged before and after blue light (488 nm) stimulation. Insets show magnified views of the boxed regions. Schematic outlines of perilipin 1 signal morphology are shown on the right. Scale bar: 4 µm. **D.** <u>Upper panel</u>: Schematic of the CRY2-based optogenetic system using an empty vector (EV), the perilipin 1 C-terminus (perilipin 1-CT^WT^), and phase separation–deficient mutant (perilipin 1-CT^FY4A^) constructs, all of which were fused to CRY2olig-mScarlet with a nuclear export signal (NES). <u>Lower</u> <u>panels</u>: differentiated brown adipocytes expressing the indicated constructs were imaged before (−2 s) and after blue light stimulation at 0, 30, and 60 seconds. Scale bar: 4 µm. **E.** Mouse perilipin 1^WT^ or the phase separation-deficient mutant perilipin 1^FY4A^ were reintroduced to *Plin1* KO preadipocytes. Differentiated adipocytes were incubated at 37 °C or 27 °C for 90 minutes, followed by norepinephrine (NE) at 0.1 μM for 10 minutes. Immunoblot analysis of perilipin 1 phosphorylation using an anti–phospho-PKA substrate antibody. **F.** Schematic of lipolysis assay in iWAT explants of perilipin 1^FY4A^ knock-in mice and littermate control mice at 37 °C or 27 °C in the presence or absence of NE. **G.** Lipolysis assays in iWAT explants of perilipin 1^FY4A^ knock-in mice and littermate controls at 37 °C. *n* = 6 explants from 2 independent mice. Data are normalized to control mice at 9 h of NE treatment, and are presented as mean ± s.e.m. Two-way ANOVA with Šídák’s multiple comparisons test. **H.** Lipolysis assays in iWAT explants at 27 °C, as shown in (**G**). **I.** *In vitro* phase separation assays using purified GFP-tagged human perilipin 1 C-terminal proteins (GFP-hPerilipin 1-CT^WT^, GFP-hPerilipin 1-CT^K520R^) at the indicated protein concentrations under the same conditions shown in (**B**). Scale bar: 4 µm. **J.** Human perilipin 1^WT^ or perilipin 1^K520R^ mutant was reintroduced into *Plin1* KO preadipocytes. Differentiated adipocytes incubated at 37 °C were treated with norepinephrine (NE) at 0.1 μM for 10 min. Immunoblot analysis shows phospho-perilipin 1^S497^. Right: Quantification, as presented by mean ± s.e.m. *n* = 4. Unpaired *t*-test. **K.** Lipolysis assessed by glycerol release following 6 h NE stimulation in adipocytes, shown in (**J**). Glycerol levels in the conditioned medium were normalized to total protein. *n* = 6 per group. Data are presented as mean ± s.e.m. Unpaired *t*-test.

The C-terminal region of perilipin 1 contains several aromatic residues (F418, F479, F495, F496, and Y510) that could plausibly act as “sticker” elements that promote multivalent cation–π interactions during phase separation (*43, 44, 55*). Thus, we generated alanine-substitution mutants targeting four key residues (FY4A) and purified the proteins (**Supplementary Figure 7E**). While the purified C-terminal domain of wild-type perilipin 1 (perilipin 1-CT^WT^) and the phosphorylation-deficient mutant (perilipin 1-CT^3SA^) displayed dose-dependent condensation at 27 °C, the FY4A mutant (perilipin 1-CT^FY4A^) failed to form condensates under the same conditions (**Figure 4B**).

To seek structural insights into perilipin 1 LLPS without altering temperature, we next used a light-activated LLPS system (*56–58*) in which the Arabidopsis photoreceptor cryptochrome-2 oligomerization domain (CRY2olig) was fused to the N-terminus of wild-type perilipin 1 (CRY2olig-perilipin 1^WT^) in adipocytes. Upon blue-light exposure (488 nm) for 30 seconds, CRY2olig-perilipin 1^WT^ rapidly formed condensates clustered on lipid droplets even at 37 °C (top panels in **Figure 4C**). In agreement with the above results, the light-induced perilipin 1 condensates required the C-terminal domain (middle panels in **Figure 4C**). Importantly, mutation of the four key aromatic residues (FY4A) abolished light-induced condensate formation (lower panels in **Figure 4C**). To test the structural sufficiency of the C-terminal domain, we then fused CRY2olig to the C-terminal region of perilipin 1 (aa 406-517, illustrated in the top panels in **Figure 4D**). Within 30 seconds of blue-light activation, the perilipin 1 C-terminus rapidly formed condensates (**Figure 4D, Supplementary Figure 9A**). Mutation of aromatic and positively charged (K/R) residues throughout this region to alanine (20 residues in total; CRY2olig-perilipin 1-CT^FY20A^) prevented condensate formation (**Supplementary Figure 9B**). Subsequently, we found that mutation of the four key aromatic residues alone (CRY2olig-perilipin 1-CT^FY4A^) was sufficient to abolish light-induced LLPS (lower panels in **Figure 4D**). This structural requirement for the aromatic residues in the C-terminus is also observed in HEK293 cells (**Supplementary Figure 9C**).

Next, we tested the biological significance of perilipin 1 LLPS by two approaches. First, wild-type perilipin 1 (Perilipin 1^WT^) or the phase-separation incompetent form of perilipin 1 (Perilipin 1^FY4A^) was reintroduced into CRISPR-*Plin1* KO cells (used in Figure 2I), which were then differentiated into mature adipocytes at 37 °C. Subsequently, these adipocytes were treated with a low dose of NE (0.1 µM) at 27 °C or 37 °C. We found that NE treatment induced perilipin 1 phosphorylation to a similar degree in perilipin 1^WT^ and perilipin 1^FY4A^ cells at 37 °C. However, cooling to 27 °C markedly potentiated NE-induced perilipin 1 phosphorylation in perilipin 1^WT^ cells, whereas this effect was significantly blunted in perilipin 1^FY4A^ cells (**Figure 4E**, quantification in **Supplementary Figure 10A**). Perilipin 1^FY4A^ cells were also resistant to cooling-enhanced lipolysis in the presence of NE (**Supplementary Figure 10B, C**). Second, we generated knock-in mice carrying the LLPS-incompetent perilipin 1 (Perilipin 1^YF4A^) (**Supplementary Figure 10D**). We then isolated the inguinal WAT from perilipin 1^YF4A^ knock-in mice and littermate wild-type control mice and performed *ex vivo* lipolysis assays at 27 °C or 37 °C (**Figure 4F**). At 37 °C, we found no significant difference in the basal and NE-induced lipolysis between knock-in mice and control mice (**Figure 4G**). However, at 27 °C, NE-induced lipolysis was significantly impaired in Perilipin 1^YF4A^ knock-in mice compared to control mice. (**Figure 4H**). These data suggest that cold-induced perilipin 1 LLPS requires four aromatic residues in the C-terminal domain, which are critical for cooling-enhanced lipolysis in vivo.

These results also raised the possibility that genetic variants within the IDR of perilipin 1 could modulate lipolysis capacity. Notably, several disease-causing mutations in the C-terminal region of perilipin 1 have been identified in humans with familial partial lipodystrophy, and some of these variants are associated with excess lipolysis rather than impaired lipogenesis (*59–61*). One intriguing example is a K520R substitution located in the C-terminus of perilipin 1 caused by the c.1559A>G missense mutation (RCV005052206.1). Because arginine can engage more strongly than lysine in cation-π interactions (*62*), we examined whether this missense variant might increase the LLPS propensity of perilipin 1. To test this, we generated purified K520R human perilipin 1 protein along with the wild-type form (**Supplementary Figure 10E**). We found that purified K520R human perilipin 1 protein formed condensates at a lower concentration than wild-type perilipin 1 in a cell-free system at 27 °C (**Figure 4I**). When wild-type or K520R perilipin 1 was reintroduced into *Plin1* KO cells and cultured at 37 °C, the K520R mutant produced significantly higher perilipin 1^S497^ phosphorylation in response to NE (**Figure 4J**, quantification on the right). This increase was accompanied by greater glycerol release and a larger reduction in intracellular lipid content (**Figure 4K, Supplementary Figure 10F**). Together, these results suggest that a lipodystrophy-associated mutation in the perilipin 1 IDR enhances its LLPS and lipolysis.

### Cooling-induced perilipin 1 phase separation increases PKACα-perilipin 1 proximity at lipid droplets

Phase separation can reorganize signaling molecules, thereby reshaping cellular signaling cascades (*43, 63–65*). Hence, we asked how cooling-induced perilipin 1 LLPS amplifies PKA signaling and lipolysis. PKA consists of a catalytic subunit (PKA-Cα) and regulatory subunits such as PKA-RII, which help localize the kinase via A-kinase anchoring proteins (AKAPs) (*66, 67*). PKA-Cα itself lacks an obvious IDR (**Supplementary Figure 11A**), and cooling did not induce condensation of purified mCherry-tagged PKA-Cα alone at a concentration (10 µM), the condition where purified perilipin 1 readily formed dynamic condensates (left panels in **Figure 5A**). However, when both proteins were mixed and incubated at 27 °C, PKA-Cα partitioned into perilipin 1 condensates and co-localized with perilipin 1. This cold-induced formation of perilipin 1-PKA-Cα condensates was blocked by 1,6-HD (right panels in **Figure 5A**). FRAP experiments also showed coincident recovery of perilipin 1 and PKA-Cα fluorescence after photobleaching (**Figure 5B**). These findings indicate that cold-induced perilipin 1 condensates increase the local proximity of PKA-Cα to perilipin 1.

**Figure 5.**
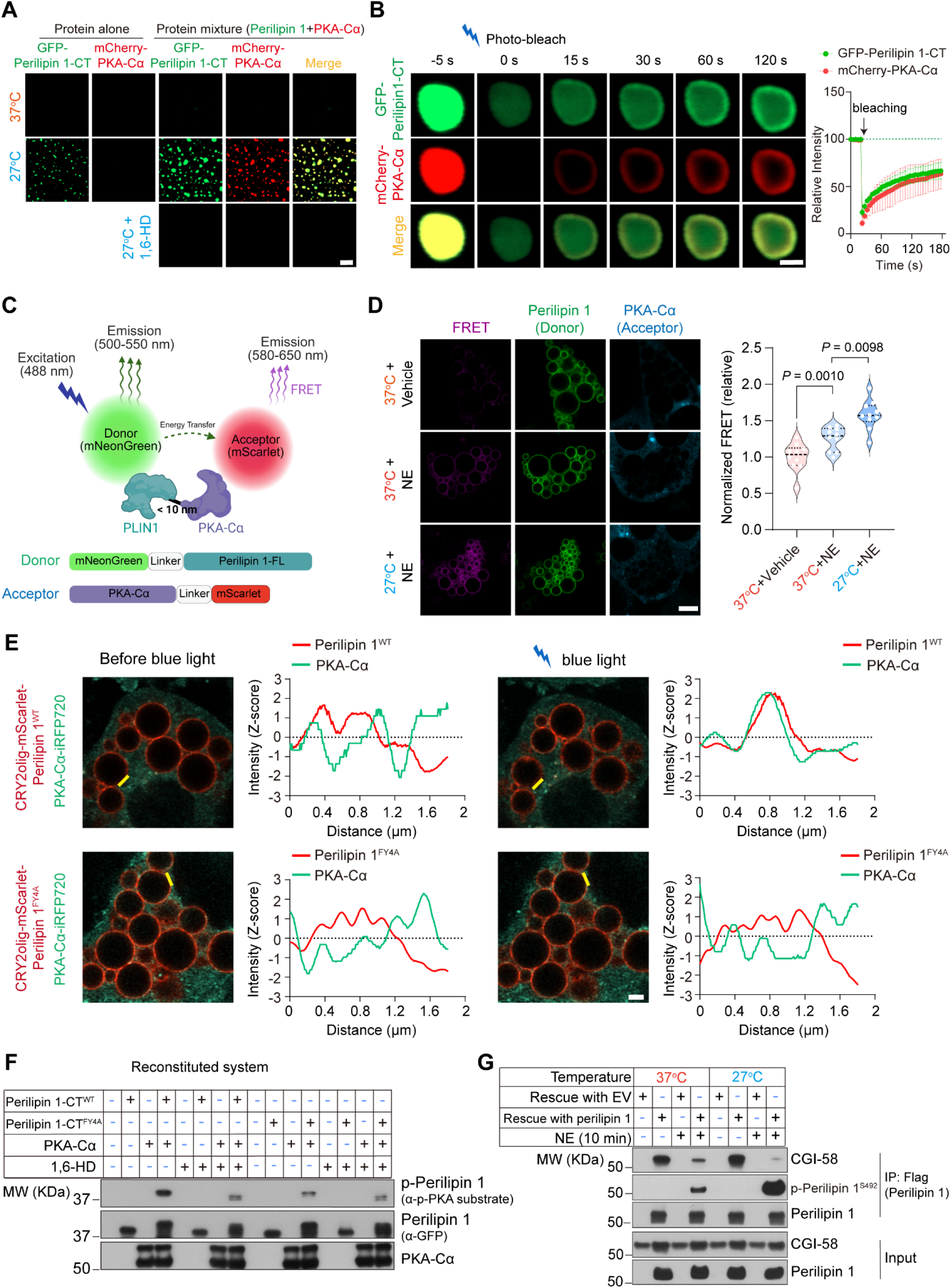
Cooling-induced perilipin 1 LLPS promotes PKA-Cα recruitment to lipid droplets. **A.** *In vitro* phase separation assays using purified GFP-perilipin 1-CT and/or mCherry-PKA-Cα proteins. Either GFP–perilipin 1-CT or mCherry-PKA-Cα alone at 10 μM was incubated in a buffer containing 150 mM NaCl, 25 mM HEPES (pH 7.4), 1 mM DTT, and 7.5% PEG-8000 at 37 °C or 27 °C. The right three columns show droplet formation when both proteins were mixed under the same conditions, in the absence or presence of 10% 1,6-HD. Scale bar: 4 µm. **B.** <u>Left:</u> Representative fluorescence recovery images after photobleaching (FRAP) of droplets formed by GFP-perilipin 1-CT and mCherry-PKA-Cα at 27 °C. Scale bar: 1 µm. <u>Right:</u> Quantification of fluorescence recovery within the bleached region. *n* = 3 independent experiments. **C.** Schematic illustration of the Förster resonance energy transfer (FRET) system using mNeonGreen– perilipin 1 as the donor and PKA-Cα–mScarlet as the acceptor. Upon excitation at 488 nm, energy transfer occurs when the two proteins are within <10 nm, leading to acceptor emission (580-650 nm). Donor emission (500-550 nm) and acceptor signals were used to quantify FRET efficiency. **D.** <u>Left</u>: Representative images of FRET-based proximity assay between perilipin 1 and PKA-Cα in adipocytes. Differentiated brown adipocytes stably expressing mNeonGreen–perilipin 1 (Donor) and PKA-Cα–mScarlet (Acceptor) were incubated at 37 °C or 27 °C for 90 min, followed by NE stimulation at 0.1 μM for 5 min. Scale bar: 5 µm. <u>Right</u>: Quantification of FRET signals. *n* = 10 adipocytes per group. Data are presented as mean ± s.e.m. One-way ANOVA followed by Tukey’s multiple comparisons test. **E.** Optogenetic clustering assay using CRY2olig-mScarlet-tagged perilipin 1 (WT or FY4A mutant) and PKA-Cα–iRFP720 in differentiated brown adipocytes. Images were captured before and after blue light (488nm, 30 seconds) stimulation to induce perilipin 1 clustering. Line-scan analysis (indicated by the yellow line) shows fluorescence intensity profiles of perilipin 1 and PKA-Cα across the lipid droplets. Upon blue light stimulation, the PKA-Cα signals were enriched at perilipin 1 puncta in cells expressing perilipin 1^WT^ but not in cells expressing perilipin 1^FY4A^. Scale bar: 2 µm. <u>Right</u>: Quantification by fluorescence intensity (Z-score) along the indicated line shown on the right side next to the image. **F.** Reconstitution of PKA-mediated phosphorylation of perilipin 1. Purified GFP-perilipin 1 C-terminus protein (GFP-perilipin 1-CT^WT^) or a phase-separation-deficient mutant (GFP-perilipin 1-CT^FY4A^) was incubated with purified mCherry-PKA-Cα at 27 °C for 20 min. Where indicated, 1,6-HD (5%) was added to disrupt LLPS. Immunoblot analysis of phospho-perilipin 1 using an anti-p-PKA substrate antibody, total perilipin 1 and PKA-Cα. **G.** Cooling-induced PKA-mediated phosphorylation of perilipin 1 promotes CGI-58 release. Co-immunoprecipitation (Co-IP) assays were performed in *Plin1* KO adipocytes expressing Flag-perilipin 1 or an empty vector (EV). Differentiated adipocytes were incubated at 37 °C or 27 °C for 90 min, followed by NE stimulation at 0.1 μM for 10 min. Flag-perilipin 1 complexes were immunoprecipitated with a Flag antibody and analyzed by immunoblotting using antibodies against Flag (Perilipin 1), perilipin 1^S492^, and CGI-58. Input (Perilipin 1 and CGI-58) was shown in the bottom panels.

We next used Förster resonance energy transfer (FRET) imaging to further examine the underlying mechanism. To this end, we developed mNeonGreen-perilipin 1 and PKA-Cα-mScarlet fusion constructs and expressed them in adipocytes (**Figure 5C**). We found that mNeonGreen-perilipin 1 signals were localized predominantly to lipid droplets; upon NE stimulation, FRET efficiency was detected on lipid droplets at 37 °C (**Figure 5D**). Notably, NE treatment at 27 °C significantly increased FRET efficiency, indicating that cooling enhances the proximity between perilipin 1 and PKA-Cα under β-adrenergic stimulation (quantification on the right in **Figure 5D**). On the other hand, this was not the case for PKA-Cα and PKA-RII: we developed a FRET system to examine whether cooling also alters their proximity (**Supplementary Figure 11B**). NE decreased the proximity between PKA-Cα and PKA-RII at 37 °C (**Supplementary Figure 11C**), consistent with cAMP-dependent release of PKA-Cα from regulatory subunits (*66, 67*). However, cooling did not further alter this interaction. These results were in agreement with our observation that cooling selectively amplified PKA-mediated perilipin 1 phosphorylation rather than broadly activating the PKA signaling pathway (see **Figure 2**).

The effect of cooling could be reflected in changes to other proteins, including AKAPs. We therefore asked whether inducing perilipin 1 condensates via the 4 aromatic residue-dependent mechanism sufficiently augments PKA recruitment to lipid droplets without cooling. To test this, we used the CRY2olig-based light-inducible system described above in which CRY2olig-perilipin 1^WT^ or CRY2olig-perilipin 1^FY4A^ was expressed together with PKA-Cα in adipocytes and examined the co-localization before and after blue-light exposure at 37 °C. Before light activation, the CRY2olig-perilipin 1 signal was distributed relatively uniformly on lipid droplets, with limited overlap with the PKA-Cα signal (green) (upper left panels in **Figure 5E**). Following 30 seconds of blue-light activation, however, CRY2olig-perilipin 1^WT^ formed punctate condensates on lipid droplets, accompanied by increased localization of PKA-Cα to these puncta (upper right panels in **Figure 5E**). In contrast, this light-induced co-localization was not observed in cells expressing CRY2olig-perilipin 1^FY4A^ (lower panels in **Figure 5E**). These results suggest that induced perilipin 1 condensation promotes the recruitment of PKA-Cα onto lipid droplets, which requires the 4 aromatic residues in the IDR of perilipin 1.

These results also prompted us to reconstitute cooling-enhanced phosphorylation of perilipin 1 by PKA in a cell-free system. To do this, we incubated purified wild-type perilipin 1 (Perilipin 1-CT^WT^) or the FY4A mutant (Perilipin 1-CT^FY4A^) proteins with purified PKA-Cα, ATP, and Mg^2+^ at 27 °C in the presence or absence of 1,6-HD, and subsequently analyzed the reaction mixtures by electrophoresis followed by immunoblotting for perilipin 1 phosphorylation (**Supplementary Figure 11D**). We observed robust phosphorylation of perilipin 1S492 in the wild-type mixture at 27 °C, whereas it was markedly reduced in the presence of 1,6-HD. By contrast, substantially less perilipin 1^S492^ phosphorylation was detected in the reaction mixture with perilipin 1-CT^FY4A^ (**Figure 5F**). Perilipin 1 binds CGI-58 (also known as ABHD5) through its C-terminal region to suppress basal lipolysis, while phosphorylation of perilipin 1 at S492 and S517 promotes CGI-58 release, enabling its interaction with ATGL and activation of lipase activity (*68, 69*). We therefore examined whether cold-inducible perilipin 1 LLPS modulates this interaction. To do so, Flag-tagged perilipin 1 was purified from adipocytes that were stimulated with NE at 37 °C or 27 °C. While perilipin 1 protein abundance was unchanged between conditions, NE induced higher perilipin 1^S492^ phosphorylation levels at 27 °C than at 37 °C. Consistent with this, the perilipin 1-CGI-58 interaction was markedly reduced at 27 °C, indicating enhanced CGI-58 release under cooled conditions (**Figure 5G**). These results indicate that cooling-induced perilipin 1 condensation increased proximity to PKA-Cα and enhanced its local recruitment to lipid droplets, leading to enhanced perilipin 1^S492^ phosphorylation, CGI-58 release, and ATGL-mediated lipolysis.

### Engineering perilipin 1 phase separation drives lipolysis

Although systemic cold exposure and β-adrenergic signaling stimulate lipolysis, they can also trigger vasoconstriction and increase blood pressure, a major risk factor for cardiovascular disease (*70, 71*). Hence, we asked whether engineered enhancement of perilipin 1 LLPS could promote lipolysis even at ambient temperature. To test this idea, we used two complementary approaches. First, CRY2olig-perilipin 1^WT^ was introduced into CRISPR-*Plin1* KO preadipocytes, which were then differentiated into adipocytes and treated with NE at 0.1 µM or vehicle at 37 °C. During this period, a subset of cells was exposed to blue light for 1 min every 30 min (total light exposure time, 12 min) (**Figure 6A**). Whereas blue light alone did not induce perilipin 1^S492^ phosphorylation, blue-light activation together with NE robustly amplified perilipin 1^S492^ phosphorylation in CRY2olig-perilipin 1^WT^-expressing adipocytes at 37 °C (**Figure 6B**). This increase in p-perilipin 1^S492^ was accompanied by enhanced lipolysis, reflected by reduced intracellular TG content (**Figure 6C**) and increased NE-stimulated glycerol release into the conditioned medium (**Figure 6D**).

**Figure 6.**
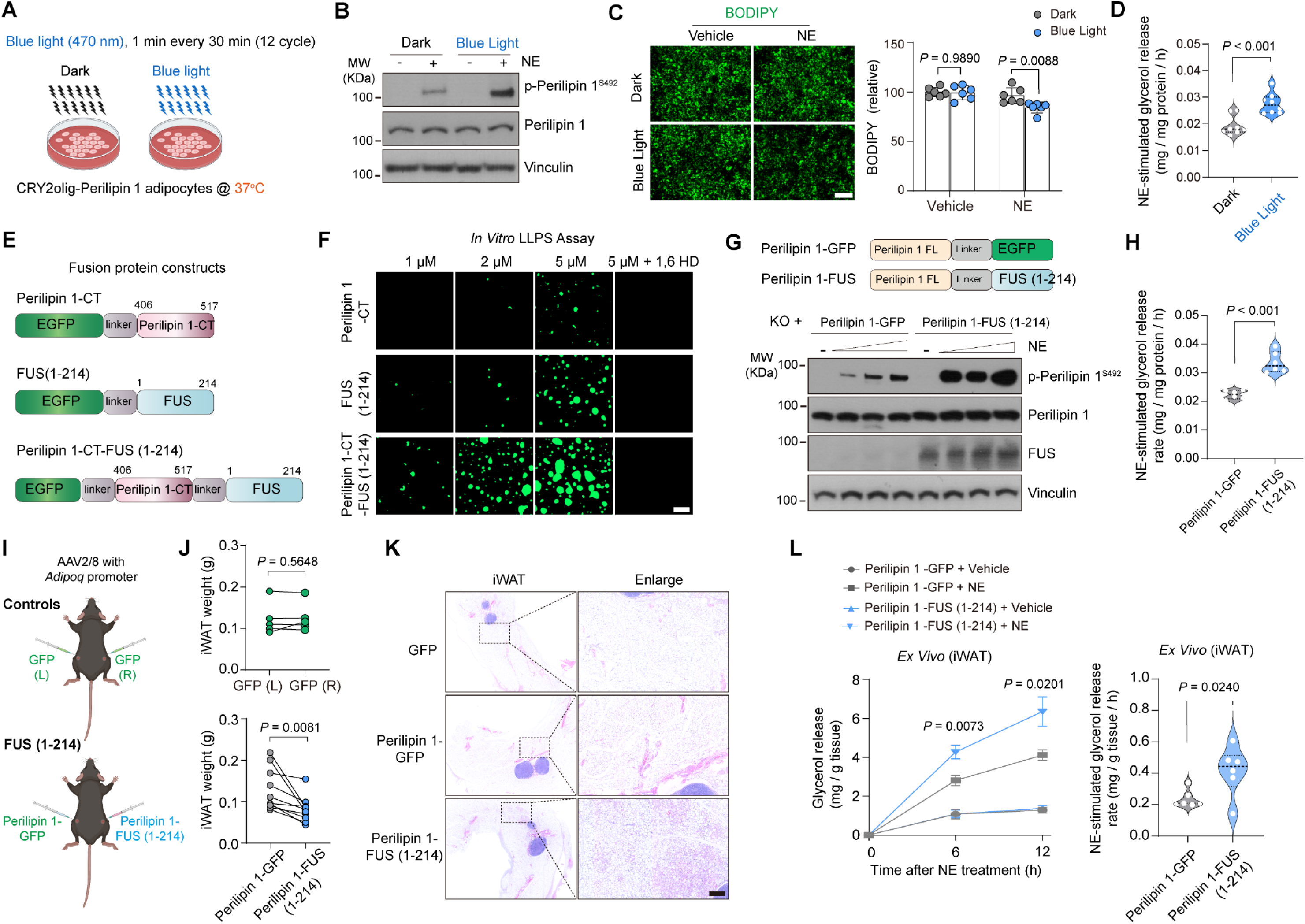
Enhanced lipolysis by engineering perilipin 1 phase separation. **A.** Schematic illustration of the optogenetic-based perilipin 1 phase separation. *Plin1* KO adipocytes expressing CRY2olig–perilipin 1 were subjected to dark or blue light (470 nm) stimulation (1 min exposure every 30 min; 12 min exposure in total) in the presence or absence of NE (0.1 μM) at 37 ^°^C. **B.** CRY2olig-perilipin 1-expressing adipocytes in (**A**) were treated with NE or vehicle for 10 min, with blue light. Immunoblot analysis shows phospho-perilipin 1^S492^, total perilipin 1, and Vinculin as controls. **C.** <u>Left:</u> Representative BODIPY staining of adipocytes under the indicated conditions. Cells were stimulated with NE for 6 h under blue light illumination. Scale bar: 130 µm. <u>Right:</u> Quantification of BODIPY fluorescence intensity. *n* = 6 per group. Data are presented as mean ± s.e.m. Two-way ANOVA with Šídák’s multiple comparisons test. **D.** Lipolysis was assessed by measuring NE-stimulated glycerol release into the conditioned medium under dark or blue light conditions in (**C**). Data are normalized to total protein level and presented as mean ± s.e.m. *n* = 8 per group. Unpaired *t*-test. **E.** Schematic of perilipin 1 constructs, including GFP–perilipin 1 C-terminus (GFP-perilipin 1-CT), GFP–FUS (1–214), and the chimeric construct GFP-perilipin 1-CT–FUS (1–214). **F.** *In vitro* phase separation assays using the purified proteins at the indicated concentrations. Representative images show enhanced droplet formation by perilipin 1-CT–FUS compared to perilipin 1-CT or FUS alone, which was abolished by 10% 1,6-HD. Scale bar: 4 µm. **G.** <u>Upper panel:</u> Schematic of perilipin 1 constructs used in the experiment. <u>Lower panel</u>: *Plin1* KO adipocytes expressing perilipin 1-GFP or perilipin 1-FUS (1–214) were stimulated with norepinephrine (NE) at 0.05, 0.1, or 0.2 μM for 10 minutes. Immunoblot analysis shows phospho-perilipin 1^S492^, total perilipin 1, FUS, and Vinculin as a control. **H.** Lipolysis was assessed by measuring glycerol release into the conditioned media in (**G**) in response to NE at 0.1 μM for 8 h. Data were normalized to total protein. *n* = 6 per group. Unpaired *t*-test. **I.** Schematic of the experimental design. AAV2/8 driven by the adiponectin (*Adipoq*) promoter was used to express GFP control, perilipin 1-GFP, or perilipin 1-FUS (1–214) in the inguinal white adipose tissue (iWAT) of mice housed at room temperature. For control mice, the GFP virus was injected bilaterally. For the FUS group, AAV-perilipin 1-GFP was injected into one side of iWAT, and AAV-perilipin 1-FUS was injected into the other side of iWAT of the same mouse. Tissues were collected 3 weeks post-injection. **J.** iWAT weight from the indicated mice in (**I**). *n* = 5 mice for control group, n= 10 mice for FUS group. Data are presented as mean ± s.e.m. Paired *t*-test. **K.** Representative H&E staining of the iWAT sections expressing GFP, perilipin 1-GFP, or perilipin 1-FUS (1–214) in (**J**). <u>Left</u>, whole-slide scans; <u>Right</u>, magnified views of the indicated regions. Scale bar: 200 μm. **L.** Lipolysis assays in *ex vivo* iWAT explants from the indicated groups. Explants were stimulated with NE at 1 μM. <u>Left</u>: Time course of glycerol release into the conditioned medium normalized to tissue weight. <u>Right</u>: NE-stimulated glycerol release rate over 12 hours. *n* = 6 explants from 2 independent mice per group. Data are presented as mean ± s.e.m. Unpaired *t*-test.

As a second strategy, we used the low-complexity domain of FUS, a well-characterized driver of LLPS that promotes condensate formation when fused to heterologous proteins (*58, 72, 73*). We fused the FUS low-complexity domain (aa 1-214) to the C-terminus of perilipin 1 to generate an LLPS-competent perilipin 1 protein (**Figure 6E, Supplementary Figure 12A**). In a cell-free reconstituted system, GFP-perilipin 1-CT-FUS (1–214) formed condensates in a dose-dependent manner at 37 °C and exhibited a much lower threshold for condensation than GFP-perilipin 1-CT or GFP-FUS (1–214) alone (**Figure 6F**). The condensate formations were blocked by 1,6-HD treatment (right panels of **Figure 6F**). We then expressed perilipin 1-FUS (1–214) or perilipin 1-GFP in *Plin1* KO preadipocytes and differentiated them into mature adipocytes. Both constructs restored lipid accumulation to a similar degree (**Supplementary Figure 12B**). Although enhanced perilipin 1 condensate formation alone did not trigger perilipin 1 phosphorylation, NE treatment led to a stronger induction of p-perilipin 1^S492^ in perilipin 1-FUS (1–214) cells than in perilipin 1-GFP cells in a dose-dependent manner (**Figure 6G**). This increase was accompanied by enhanced NE-induced lipolysis, as measured by glycerol release into the medium (**Figure 6H**). We note that overexpression of Flag-FUS (1–214) alone did not affect NE-induced perilipin 1^S492^ phosphorylation (**Supplementary Figure 12C**), indicating that the effect was not due to the action of FUS *per se*.

Finally, we examined whether enhanced perilipin 1 phase separation alters adipose tissue mass *in vivo* by ectopically delivering the LLPS-competent perilipin 1 into inguinal WAT. To this end, we generated AAV2/8 vectors expressing perilipin 1-FUS (1–214), perilipin 1-GFP, or GFP alone under the adipocyte-specific *Adipoq* promoter (**Supplementary Figure 12D**). Perilipin 1-FUS (1–214) or perilipin 1-GFP were delivered to contralateral inguinal WAT depots in the same mice, enabling a paired comparison independent of systemic metabolism, such as food intake and circulating hormones (**Figure 6I**). Mice were maintained at room temperature, where sympathetic tone remained active, as our results found that perilipin 1 phase separation induced lipolysis in the presence of NE. After 3 weeks, transgene expression was confirmed in adipose tissue (**Supplementary Figure 12E**). We found that perilipin 1-FUS (1–214) expression significantly reduced inguinal WAT mass relative to perilipin 1-GFP expression (**Figure 6J**). By contrast, no difference in depot mass was observed when GFP alone was delivered bilaterally as a control. The effect of perilipin 1-FUS was specific to the injected inguinal depot, as no change was detected in interscapular BAT or epididymal WAT that did not receive the AAVs (**Supplementary Figure 12F-H**). Histologically, ectopic expression of perilipin 1-FUS (1–214) reduced adipocyte size throughout the inguinal WAT depot relative to perilipin 1-GFP or GFP alone (**Figure 6K, Supplementary Figure 12I**). Furthermore, the reduction in the inguinal WAT depot mass was accompanied by elevated lipolysis, as *ex vivo* assays showed time-dependent enhancement of NE-induced glycerol release from inguinal WAT expressing perilipin 1-FUS (1–214) compared with those expressing perilipin 1-GFP (**Figure 6L**). Together, these results show that enhancing perilipin 1 phase separation is sufficient to increase NE-stimulated lipolysis and reduce local adipose mass *in vivo*.

## DISCUSSION

A prevailing principle in biology is that lower temperatures globally reduce most biochemical processes. Adipocytes emerge as a notable exception: within a physiologically relevant cool range (27-31 °C), they retain active adrenergic lipolysis. Unexpectedly, this response does not require canonical TRP-channel-mediated cold sensing. Instead, mild cooling selectively amplifies the adrenergic PKA signaling on perilipin 1 phosphorylation and its downstream lipolytic cascade. The present study shows that cooling promotes LLPS of perilipin 1, one of the most abundant and dominant lipid-droplet coat proteins in adipocytes. LLPS of this abundant protein substantially impacts lipid droplet accessibility, as cooling-induced LLPS of perilipin 1 promotes the local recruitment of PKA-Cα to lipid droplets, thereby amplifying PKA-mediated perilipin 1 phosphorylation and enhancing lipolysis (**Figure 7**).

**Figure 7.**
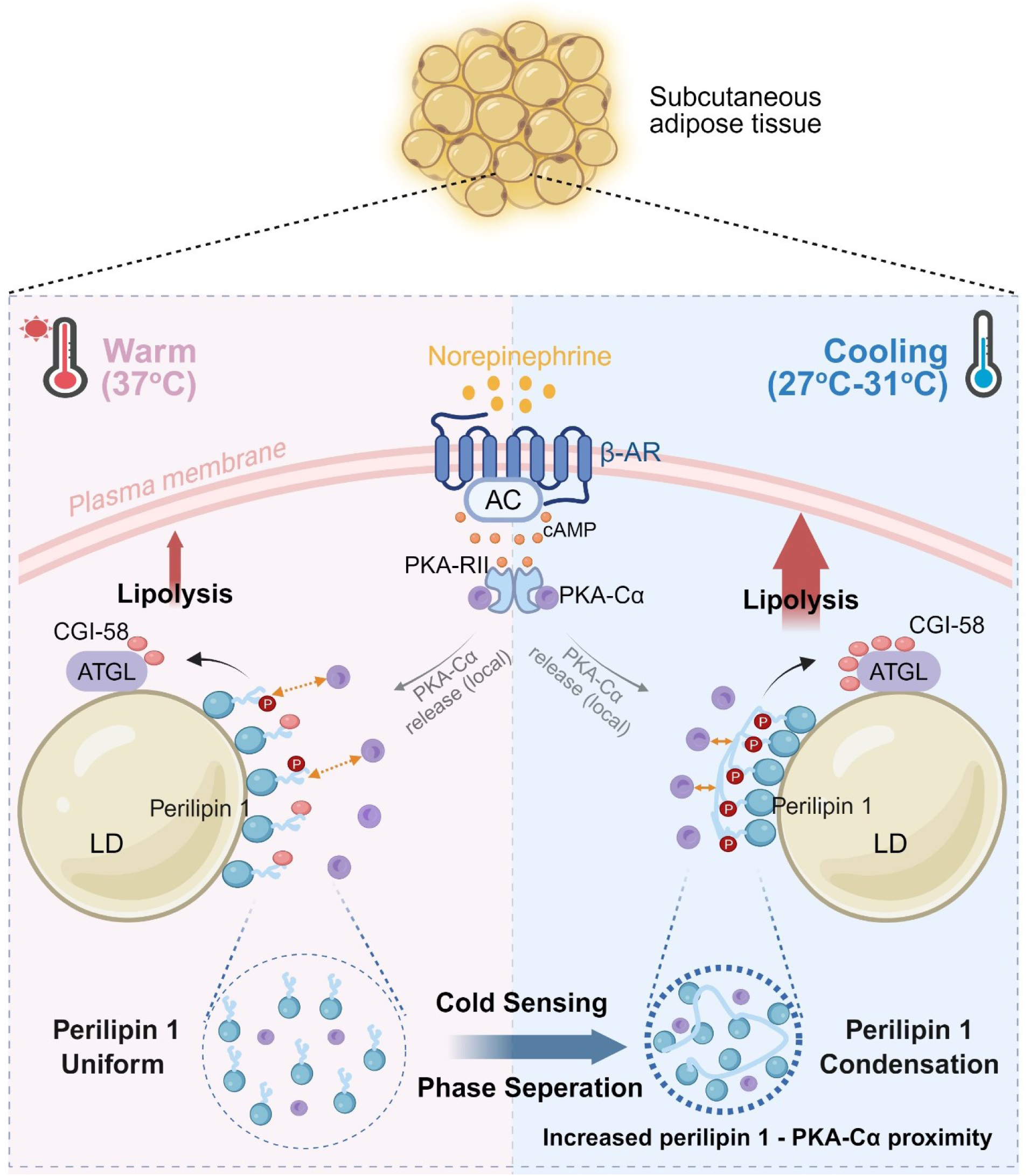
A proposed cell-autonomous mechanism by which physiologically relevant cooling temperatures promote lipolysis via perilipin 1 phase separation. Adipocytes possess a cell-intrinsic mechanism to sense physiologically relevant cooling temperatures (27– 31 °C) by inducing liquid-liquid phase separation (LLPS) of perilipin 1, the dominant lipid-coating protein. Cooling-induced LLPS of perilipin 1 promotes its condensation and the local recruitment of PKA-Cα to lipid droplet surfaces. This spatial reorganization selectively amplified PKA-mediated perilipin 1 phosphorylation, CGI-58 release, and ATGL-dependent lipolysis. The four aromatic amino acids (F/Y) in the C-terminal region are required and sufficient for cooling-induced LLPS of perilipin 1.

Whereas temperature-sensitive LLPS is recognized as a property of intrinsically disordered proteins (*43, 46, 48, 74*), its directionality is not uniform. Depending on the IDR sequence composition and solvent-mediated interactions, proteins can undergo phase separation in response to increased temperature (*a.k.a.*, a lower critical solution temperature, LCST) or decreased temperature (an upper critical solution temperature, UCST) or even exhibit a dual response across different temperature ranges. For example, a recent study on heat shock factor 1 (HSF1) reported that its phase separation is promoted by heating rather than cooling (*74*). In contrast, we identify perilipin 1 as a distinct example of cold-responsive LLPS with UCST-like behavior that occurs at 31-27 °C, well within the temperature range of subcutaneous adipose tissues, whereas such a change was not observed in PKA-Cα. Importantly, this reorganization enhances NE-induced lipolysis, suggesting that subcutaneous adipose tissue - a depot affected by outdoor temperatures - directly decodes modest local cooling to sustain lipid mobilization during chronic cold adaptation, even when other metabolic pathways (e.g., insulin signaling) are attenuated. Notably, the C-terminal IDR features required for perilipin 1 LLPS are well conserved in endotherms, whereas ectothermic perilipin 1 orthologs lack these sequence elements, implying the possibility that this thermosensitive module might evolve to support endothermic physiology. Although speculative, it is conceivable that the phase-transition temperatures of some condensates are tuned to the thermal niche experienced by a given tissue or species. Together, the present study provides a framework for understanding how subtle changes in peripheral tissue temperature influence metabolic homeostasis.

This study also led to the concept of adipose tissue-selective “cold mimetics” that facilitate lipolysis in a targeted adipose depot. We showed that both the light-inducible CRY2 system and the FUS-fusion approach enhanced PKA-mediated perilipin 1 phosphorylation and lipolysis, and targeted delivery of a perilipin 1-FUS construct to inguinal WAT increased local lipolysis and reduced depot mass *in vivo*. We note that activation of adipose lipolysis alone is insufficient to improve systemic metabolic health, because excessive lipid mobilization without coordinated oxidation can lead to ectopic lipid deposition, including hepatic steatosis (*75, 76*). In agreement, the lipodystrophy-associated perilipin 1^K520R^ variant enhanced perilipin 1 condensation and lipolytic activity. Nonetheless, the present study offers a strategy to enhance lipolytic capacity in a targeted adipose depot by delivering an LLPS-competent perilipin 1 variant or by local adipose tissue cooling. Such targeted interventions might be effective, particularly in a subset of obese or aged populations that exhibit catecholamine resistance in adipose tissue and impaired lipolysis capacity (*77–81*).

## Materials and Methods

### Animals

All animal experiments were approved by the Institutional Animal Care and Use Committee (IACUC; protocol #028-2022-25) at Beth Israel Deaconess Medical Center. Mice were housed under a 12-hour light–dark cycle with ad libitum access to food and water at 22 °C and about 45% humidity. ATGL floxed mice (Jackson Laboratory, strain #024278), wild-type C57BL/6J mice (Jackson Laboratory, strain #000664), and perilipin 1^FY4A^ knock-in mice were used in this study. Perilipin 1^FY4A^ knock-in mice, carrying the F479A, F495A, F496A, and Y510A substitutions, were generated on a C57BL/6J background by CRISPR–Cas9-mediated genome editing at the Beth Israel Deaconess Medical Center Transgenic Core. The guide RNA target sequences, donor-template sequences, and genotyping primers used in the study are listed in **Table S2**.

#### Local cold exposure model

The experimental group mice were housed at room temperature and shaved on both inguinal regions. Under anesthesia, mice were placed on either a cold pad (0-6 °C) or a warm pad (35-37 °C) for 10 min. The control group received warm-pad treatment on both inguinal regions. Skin surface temperature was monitored every 2–3 min using a thermometer and maintained above 10 °C. After treatment, mice were allowed to recover on a warm pad in a lateral position until fully awake. This procedure was repeated four times per day (total 40 min per day) for 5 consecutive days.

#### Blood glucose and serum biochemical analyses

Blood glucose levels were measured using a FreeStyle glucometer. Serum insulin, aspartate aminotransferase (AST), and alanine aminotransferase (ALT) levels were measured using a Rat/Mouse Insulin ELISA Kit (Sigma-Aldrich), an Aspartate Aminotransferase Colorimetric Activity Assay Kit (Cayman Chemical), and an Alanine Transaminase Colorimetric Activity Assay Kit (Cayman Chemical), respectively, according to the manufacturers’ instructions.

### Cells

Stromal vascular fraction (SVF) preadipocytes were isolated from inguinal white adipose tissue (iWAT) or brown adipose tissue of wild-type C57BL/6J mice or ATGL floxed mice. To generate immortalized preadipocytes, SVF cells were infected with retrovirus expressing SV40 large T antigen as previously described (*82–84*). TRPM8-knockout and perilipin 1-knockout preadipocytes were generated by infecting immortalized brown preadipocytes with eSpCas9-LentiCRISPR v2 lentivirus expressing either a scrambled control sgRNA (sgCon) or sgRNAs targeting *Trpm8* (sg*Trpm8*-A, -B, or -C) or sgRNAs targeting *Plin1* (sg *Plin1* -A or -B). For re-expression experiments, *Plin1* -KO preadipocytes (sg *Plin1* -A) were infected with retrovirus expressing mouse Flag-perilipin 1, mouse perilipin 1^WT^, mouse perilipin 1^FY4A^, mouse perilipin 1-GFP, mouse perilipin 1–FUS (1–214), human perilipin 1^WT^, or human perilipin 1^K520R^, followed by selection with blasticidin (10 μg ml⁻¹). Unless otherwise specified, cells were maintained in high-glucose DMEM supplemented with 10% fetal bovine serum (FBS), 1% penicillin–streptomycin, and 1% GlutaMAX at 37 °C with 5% CO₂. White adipocyte differentiation was induced in high-glucose DMEM supplemented with 10% FBS, 1% penicillin–streptomycin, 1% GlutaMAX, insulin (5 μg ml⁻¹), triiodothyronine (T3, 1 nM), rosiglitazone (1 μM), isobutylmethylxanthine (IBMX, 0.5 mM), indomethacin (125 nM), and dexamethasone (1 μg ml⁻¹). After 48 h, cells were maintained in medium containing insulin (5 μg ml⁻¹), T3 (1 nM), and rosiglitazone (1 μM) for an additional 4 days. Brown adipocyte differentiation was induced in high-glucose DMEM supplemented with 10% FBS, 1% penicillin–streptomycin, 1% GlutaMAX, insulin (0.5 μg ml⁻¹), T3 (1 nM), IBMX (0.5 mM), indomethacin (125 nM), and dexamethasone (1 μg ml⁻¹). After 48 h, cells were maintained in medium containing insulin (0.5 μg ml⁻¹) and T3 (1 nM) for an additional 4 days.

#### Human primary adipocyte culture

Primary human subcutaneous adipose progenitor cells were obtained from the laboratory of Evan Rosen. All procedures were approved by the Institutional Review Board (IRB #: 2011P000079) at Beth Israel Deaconess Medical Center, and all samples were de-identified. Cells were derived from subcutaneous adipose tissue of two female donors: a 68-year-old (body mass index: 29 kg m^-2^, thigh depot) and a 48-year-old (body mass index: 31 kg m^-2^, abdominal depot). Cells were maintained and expanded in EGM-2MV medium (CC-4147, Lonza). Adipogenic differentiation was induced upon confluence by culturing cells in high-glucose DMEM supplemented with 10% FBS and 1% penicillin– streptomycin, in the presence of 0.5 mM IBMX, 0.25 μM dexamethasone, 5 μg ml⁻¹ insulin, and 1 μM rosiglitazone for 3 days. Cells were subsequently maintained in high-glucose DMEM containing 10% FBS, with medium replaced every 2–3 days until experiments.

### DNA constructs, Viruses and Antibodies

Mouse and human perilipin 1 wild-type, mutant, and truncated cDNAs were synthesized (Integrated DNA Technologies, IDT) and cloned into a blasticidin-resistant pMSCV vector (Addgene, #75085) or pLenti-CMV vector (Addgene, #125133). Mouse perilipin 2, perilipin 3, and perilipin 5 cDNAs were synthesized (IDT) and cloned into pLenti-CMV vector. Mouse PKA-Cα and PKA-RII cDNAs were synthesized (IDT) and cloned into pLenti-CMV vector. The CRY2olig construct fused to mScarlet was synthesized (IDT) and cloned into the pMSCV vector. sgRNAs targeting mouse *Trpm8*, mouse *Plin1*, or a scrambled control (sgCon) were cloned into an eSpCas9-LentiCRISPR v2 backbone in which the puromycin resistance cassette was replaced with a blasticidin resistance cassette. The original eSpCas9-LentiCRISPR v2 plasmid was obtained from GenScript. All constructs were verified by Sanger sequencing. sgRNA sequences are provided in **Table S3.**

#### Virus production and infection

Retrovirus and lentivirus were produced in Lenti-X 293T packaging cells by transfection with 10 μg of transfer plasmids and 10 μg of packaging plasmids using GenJet DNA Transfection Reagent (SignaGen Laboratories). For retrovirus production, gag-pol and pMD2.G were used, whereas psPAX2 and pMD2.G were used for lentivirus production.

*Antibodies*: anti-Perilipin 1 (9349S, Cell Signaling Technology), anti-Perilipin 1 (27716-1-AP, Proteintech), anti-Perilipin 2 (15294-1-AP, Proteintech), anti-Perilipin 3 (10694-1-AP, Proteintech), anti-Perilipin 4 (55404-1-AP, Proteintech), anti-Perilipin 5 (26951-1-AP, Proteintech), Anti-P-Perilipin 1-Serine 497 (4855, Vala Sciences), anti-ATGL (2138S, Cell Signaling Technology), anti-PKA-Cα (4782S, Cell Signaling Technology), anti-P-AKT-Serine 473 (9271S, Cell Signaling Technology), anti-AKT (4691S, Cell Signaling Technology), anti-P-SMAD2-Serine 465/467 (3108S, Cell Signaling Technology), anti-SMAD2/3 (5678S, Cell Signaling Technology), anti-P-PKA substrate (9621S, Cell Signaling Technology), anti-FUS (67840S, Cell Signaling Technology), anti-GFP (sc-53882, Santa Cruz), anti-Vinculin (66305-1-IG, Proteintech), anti-Flag (8146S, Cell Signaling Technology), anti-β-actin (A3854, Sigma-Aldrich), goat anti-rabbit light chain HRP-conjugated antibody (NBP2-75935, Novus Biologicals), goat anti-mouse light chain HRP-conjugated antibody (91196S, Cell Signaling Technology), and Alexa Fluor 555 (ab150114, Abcam).

#### AAV-mediated gene expression in vivo

AAV2/8 vectors driven by the adiponectin (*Adipoq*) promoter were used to express GFP, GFP-perilipin 1, perilipin 1–GFP, or perilipin 1–FUS (1–214) in inguinal white adipose tissue (iWAT). For control mice, AAV-GFP was injected bilaterally. For the experimental FUS group, perilipin 1–GFP was injected into the left iWAT and perilipin 1–FUS (1–214) into the right iWAT of the same mouse. AAV (5 × 10^10^ genome copies per fat pad) was delivered via subcutaneous injection at five sites per iWAT to ensure broad coverage. Tissues were harvested 3 weeks post-injection. Mice were maintained with shaved inguinal regions throughout the expression period to facilitate injection and minimize local insulation. All AAVs were custom-generated by OBIO Technology.

### Protein expression and purification

A modified pET28a vector containing an MBP-super TEV protetease backbone (Addgene, 171782) was used for protein expression. The MBP-super TEV protetease was replaced with a Twin-Strep tag followed by SUMO1 (amino acids 1–101) by standard molecular cloning. Codon-optimized sequences encoding mCherry–PKA-Cα, GFP–perilipin 1-CT, GFP–FUS (1–214), or GFP–perilipin 1-CT–FUS (1–214) were synthesized (IDT) and cloned downstream of the SUMO1 tag. All constructs were verified by Sanger sequencing. Plasmids were transformed into E. coli C43 (DE3) cells (Thermo Fisher Scientific). Protein expression was induced at an optical density (OD₆₀₀) of ∼0.45 with 0.1 mM IPTG at 16 °C for 14–16 h. Cells were harvested and lysed by sonication in PBS supplemented with EDTA-free protease inhibitor (Roche). Cleared lysates were incubated with Strep-Tactin beads (IBA Lifesciences) at 4 °C for 6 h. Beads were washed three times with IBA wash buffer, followed by additional washes with buffer containing 150 mM NaCl and 50 mM Tris-HCl PH 7.4. Bound proteins were eluted and subjected to SUMO tag cleavage using ULP1 protease at 4 °C for 4 h. Proteins were further purified by size-exclusion chromatography (Cytiva) using a buffer containing 150 mM NaCl and 25 mM HEPES PH 7.4 and immediately used for downstream assays, including *in vitro* phase separation and kinase assays.

### *In Vitro* phase separation assay

*In vitro* phase separation assays were performed as previously described with minor modifications(*49*). Purified proteins were mixed at the indicated concentrations in LLPS buffer (150 mM NaCl, 25 mM HEPES PH 7.4, 1 mM DTT) supplemented with 7.5% PEG-8000 in a total volume of 30 μl. Where indicated, 1,6-hexanediol (10%) was included during sample preparation together with PEG-8000. Samples were incubated at the indicated temperatures for 10 min prior to imaging (Zeiss LSM900 confocal microscope).

### *In vitro* kinase assay

Purified GFP–perilipin 1-CT^WT^ or GFP–perilipin 1-CT^FY4A^ (5 μM) was incubated with purified mCherry–PKA-Cα (0.5 μM) in kinase buffer containing 150 mM NaCl, 25 mM HEPES PH 7.4, 20 mM MgCl_2_, and 200 μM ATP. Where indicated, 1,6-hexanediol (5%) was included in the reaction mixture. Reactions were carried out at 27 °C for 20 min and terminated by addition of SDS sample buffer, followed by SDS–PAGE analysis.

### Fluorescence recovery after photobleaching (FRAP) assay

FRAP assays were performed on *in vitro* phase-separated droplets or condensates formed in cells following cooling. Photobleaching was carried out using a 488 nm or 561 nm laser at 100% power on selected regions of interest (ROIs) within individual droplets or condensates. Approximately 10–30 iterations were applied to achieve sufficient bleaching. Fluorescence recovery was monitored at 5–10 s intervals. Fluorescence intensities were quantified using ImageJ and normalized to pre-bleach levels after background subtraction and correction for acquisition photobleaching. All imaging was performed using a Zeiss LSM900 confocal microscope.

### Förster resonance energy transfer (FRET) assay

Immortalized brown preadipocytes were engineered to stably express mNeonGreen–Perilipin 1 together with PKA-Cα–mScarlet or PKA-RII–mNeonGreen together with PKA-Cα–mScarlet. Cells were differentiated into mature adipocytes for 4 days prior to experiments. For temperature treatment, adipocytes were incubated at 37 °C or 27 °C for 90 min, followed by treatment with norepinephrine (NE, 0.1 μM) for 1-5 min prior to imaging. FRET imaging was performed on a Zeiss LSM900 confocal microscope using sequential frame-by-frame scanning to minimize spectral cross-talk. Donor excitation was performed at 488 nm, and emission was collected at 500–550 nm (donor channel) and 580–650 nm (FRET channel). Acceptor excitation was performed at 561 nm, and emission was collected at 580–650 nm (acceptor channel). Donor-only and acceptor-only control cells were imaged under identical settings to determine spectral bleed-through coefficients. Normalized FRET (nFRET) was calculated after correction for donor bleed-through and acceptor cross-excitation following previous study (*85*).

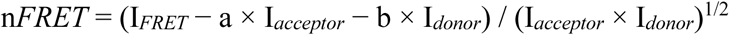

Where I_FRET_ is the raw FRET signal, I_donor_ and I_acceptor_ are the donor and acceptor intensities, and a and b represent the bleed-through coefficients determined from acceptor-only and donor-only controls, respectively.

### Optogenetic assay

Optogenetic assays were performed based on previously described CRY2 oligomerization systems with minor modifications (*58*). The mCherry tag was replaced with mScarlet to generate the CRY2olig–mScarlet construct. Protein of interest were cloned downstream of CRY2olig– mScarlet to assess their phase separation capacity. Stable adipocyte cell lines expressing the indicated fusion proteins were generated by retroviral transduction. Live-cell imaging was performed at 37 °C using a Zeiss LSM900 confocal microscope. For photoactivation, regions of interest were illuminated with a 488 nm laser at 0.3–0.6% power for approximately 10–30 iterations to induce CRY2 oligomerization and promote condensate formation. Fluorescence signals were subsequently monitored in the 561 nm channel at 5–10 s intervals. Images were acquired under identical settings across conditions.

### Lipid droplet isolation and cooling-induced condensation

Lipid droplets were isolated from cultured adipocytes or mouse iWAT as previously described with minor modifications (*86*). All procedures, prior to the indicated incubation temperature, were performed rapidly using pre-warmed reagents and equipment, with samples maintained as close to 37°C as possible. For cultured adipocytes, differentiated adipocytes were washed once with pre-warmed PBS and twice with pre-warmed LD isolation buffer (250 mM sucrose, 5 mM HEPES, and EDTA-free protease inhibitor cocktail). Cells were collected in 1.5 mL microcentrifuge tubes and homogenized by repeated pipetting with a P200 pipette without a tip. Homogenates were centrifuged at 2,000 *g* for 5 min, and the upper LD layer was collected and placed onto glass-bottom dishes. Samples were incubated at 37°C or 27°C for 10 min to allow condensate formation and then immediately imaged. For LDs isolated from iWAT, freshly dissected tissue was rapidly washed once with pre-warmed PBS and twice with pre-warmed LD isolation buffer. The tissue was minced with scissors, homogenized thoroughly with a Dounce homogenizer, and rapidly transferred into 1.5 mL microcentrifuge tubes using a P200 pipette with the tip removed. Homogenates were centrifuged at 2,000 *g* for 5 min. The upper LD layer was then collected, placed onto glass-bottom dishes, and incubated at the indicated temperature before immediate image acquisition. GFP signal distribution on LDs was quantified using Fiji software (*87*).

### Lipolysis Assays

Fully differentiated adipocytes were incubated in lipolysis medium (high-glucose DMEM supplemented with 2% fatty acid–free BSA and 1% penicillin–streptomycin) at 27 °C or 31 °C with 5% CO2 for 1 h, followed by treatment with norepinephrine (NE) for cells of mouse origin or isoproterenol for cells of human origin at the indicated concentrations for 6–10 h. We used ISO for human cells because inhibitory *α*_2_AR receptors are abundant in human adipocytes (*88*). Cells and conditioned media were collected for analysis. Glycerol and free fatty acid (FFA) levels in the culture medium were measured using a glycerol assay kit (Sigma-Aldrich) and a free fatty acid assay kit (RayBiotech), respectively, according to the manufacturers’ instructions. Values were normalized to total cellular protein content. Lipolytic activity was expressed as the rate of glycerol or FFA release over the indicated time period.

For optogenetic stimulation of lipolysis, *Plin1*-knockout adipocytes were reconstituted with CRY2olig– perilipin 1 and differentiated into mature adipocytes. Cells were incubated in lipolysis medium at 37 °C in the presence or absence of norepinephrine (NE, 0.1 μM). Blue light stimulation was applied using a 470 nm LED in 12 cycles of 1 min illumination every 30 min (total illumination time of 12 min) over a 6 h period. Conditioned media were collected for analysis of lipolytic activity as described above.

For adipose tissue explant assays, paired iWAT depots expressing perilipin 1–GFP or perilipin 1–FUS from the same mouse, as well as iWAT depots from perilipin 1^FY4A^ knock-in mice, were dissected, washed three times with PBS, and cut into ∼2 mm pieces, following a previous study (*89*). Tissue explants were pre-incubated in lipolysis medium at 37 °C for 1 h, followed by stimulation with NE (1 μM). Conditioned media were collected at indicated time point, and glycerol levels were measured as described above. Values were normalized to tissue weight. Lipolytic activity was presented as glycerol release at each time point or as NE-stimulated glycerol release rates over 12 h.

### Co-immunoprecipitation (co-IP)

*Plin1*-knockout adipocytes were reconstituted with empty vector (EV) or Flag-tagged Perilipin 1. After differentiation for 6 days, cells were incubated at 37 °C or 27 °C for 90 min, followed by treatment with or without norepinephrine (NE, 0.1 μM) for 10 min. Cells were lysed in IP buffer (150 mM NaCl, 50 mM Tris-HCl PH 7.4, 0.5% NP-40, supplemented with EDTA-free protease inhibitor) on ice. Cleared lysates were incubated with anti-Flag magnetic beads (Thermo Fisher Scientific) at 4 °C for 6 h with rotation. Beads were washed five times with wash buffer (150 mM NaCl, 50 mM Tris-HCl PH 7.4, 0.5% Triton X-100, supplemented with EDTA-free protease inhibitor) to remove non-specific binding. Immunoprecipitated proteins were analyzed by immunoblotting to assess the interaction between perilipin 1 and CGI-58.

### Metabolomics

Metabolomic sample preparation was performed following established protocols (*90*). Equal numbers of immortalized brown preadipocytes were differentiated for 6 days and subsequently incubated at 37 °C or 27 °C for 1 h, followed by treatment with norepinephrine (NE, 0.1 μM) for 10 h. Cells were rapidly washed and metabolites were extracted by addition of 300 μl pre-chilled 100% methanol (−20°C) on dry ice. Cells were scraped and lysed by vigorous shaking for 5 min. An aliquot of 200 μl extract was mixed with 200 μl chloroform and 100 μl water, vortexed thoroughly, and centrifuged at 16,000 × g. The upper aqueous phase was collected, and 150 μl was further filtered using a 10 kDa cutoff filter (Sigma-Aldrich) to remove proteins and debris. Filtered samples were transferred to LC–MS vials for analysis. Metabolite detection and data processing were performed as previously described (*91*). Metabolomics data were acquired using a UHPLC system (Vanquish Horizon, Thermo Fisher Scientific) coupled to an Orbitrap mass spectrometer (Exploris 240, Thermo Fisher Scientific). Chromatographic separation was performed using a Waters ACQUITY UPLC BEH Amide column (1.7 μm, 100 × 2.1 mm) at 25 °C. The mobile phases consisted of A (25 mM ammonium acetate and 25 mM ammonium hydroxide in water) and B (acetonitrile), operated in negative ion mode. The gradient was as follows: 95% B (0–1 min), 95–65% B (1–7 min), 65– 40% B (7–8 min), 40% B (8–9 min), 40–95% B (9–9.1 min), followed by re-equilibration at 95% B for 5.9 min. The flow rate was 0.4 ml min⁻¹. The injection volume was 2 μl. Electrospray ionization (ESI) parameters were set as follows: spray voltage, 3,500 V (positive mode) or −2,800 V (negative mode); vaporizer temperature, 350 °C; sheath gas, 50 arb; auxiliary gas, 10 arb; ion transfer tube temperature, 325 °C. Full MS scans were acquired at a resolution of 60,000 with a scan range of 70–1,050 m/z and a maximum injection time of 100 ms. Data-dependent MS/MS (ddMS2) scans were acquired at a resolution of 30,000 with a maximum injection time of 60 ms, topN = 6, isolation window of 1.0 m/z, and normalized collision energy of 30%. Dynamic exclusion was enabled. Metabolite peak areas were quantified using TraceFinder software (Thermo Fisher Scientific).

### Lipidomic

Lipidomic analysis was performed following established protocols (*91*). Equal numbers of fully differentiated adipocytes were used for each condition and subjected to temperature and norepinephrine (NE) treatments as described above. For cellular lipid extraction, lipids were isolated using a modified Bligh and Dyer method. Briefly, cell lysates were mixed with methanol:chloroform:water (2:1:0.8, v/v/v) and vortexed thoroughly. Phase separation was induced by adding additional chloroform and water to achieve a final ratio of methanol:chloroform:water (2:2:1.8, v/v/v). After centrifugation, the lower organic phase was collected, dried under vacuum for 20 min, and reconstituted in isopropanol:acetonitrile (1:1, v/v). Samples were vortexed, centrifuged for 10 min, and the supernatant was used for hydrophobic lipidomic analysis. For conditioned media, lipids were extracted using a protein precipitation–based method extraction. Briefly, media samples were mixed with pre-chilled 100% methanol (−20 °C) at a ratio of 1:8 (v/v) to achieve a final methanol concentration of 80%, followed by incubation at −80 °C for 2 h. Samples were centrifuged at 16,000 × g for 15 min, and the supernatant was collected for downstream hydrophilic lipidomic analysis. Lipidomic data acquisition and processing were performed by LipidServe according to standard procedures.

### RNA-seq and analysis

Immortalized brown adipocytes were fully differentiated and pre-incubated at either 37 °C or 27 °C for 1 h, followed by stimulation with norepinephrine (0.1 μM) for 3 h. Cells were lysed and preserved in DNA/RNA Shield (Zymo Research) without prior RNA purification, and lysates were submitted to a commercial sequencing service provider (Plasmidsaurus) for library preparation, sequencing, and primary data processing. Sequencing reads were aligned to the mouse reference genome (mm10), and gene-level expression was quantified as counts per million (CPM). For visualization, expression values were log2-transformed [log2(CPM + 1)].

### Phosphoproteomics

Fully differentiated brown adipocytes were pre-incubated at 27 °C for 90 min, followed by stimulation with norepinephrine (NE, 0.1 μM) or PBS for 10 min. Cells were rapidly washed twice with ice-cold PBS supplemented with protease inhibitors and phosphatase inhibitor cocktails 2/3 (Sigma-Aldrich), and immediately snap-frozen in liquid nitrogen for subsequent analysis. Cells were lysed in 2% SDS followed by immediate heat denaturation at 95 °C for 5 min. Lysates were further homogenized by probe sonication on ice. Protein concentration was determined using a BCA assay, and equal amounts of protein (500 µg) were taken for downstream processing. Proteins were reduced with 10 mM tris(2-carboxyethyl) phosphine (TCEP, Thermo Fisher Scientific) at 37 °C for 30 min and alkylated with 20 mM iodoacetamide (IAA, Sigma-Aldrich) at room temperature for 30 min in the dark. Proteins were subsequently precipitated using a methanol–chloroform precipitation method. The resulting protein pellets were washed with methanol, air-dried, and resuspended in 200 mM EPPS buffer. Proteins were digested overnight at 37 °C using Trypsin (Promega) and Lys-C (Wako Chemicals) at a 1:100 (w/w) enzyme-to-substrate ratio. Peptide concentrations were determined by BCA assay, and 200 µg of peptides were used for downstream labeling. Peptides were adjusted to a uniform volume with EPPS buffer and acetonitrile (ACN) was added to achieve a final concentration of 30% (v/v). TMT reagents (ThermoFisher Scientific) were added at a ratio of 3:1 (TMT:peptide, w/w), and the reaction was incubated at room temperature for 1 h in the dark. Labeling efficiency was evaluated by taking small aliquots from each sample, combining them, and analyzing by LC–MS. Following confirmation of labeling efficiency, reactions were quenched by addition of hydroxylamine and incubated for 15 min at room temperature. Combined samples were desalted using reversed-phase C18 solid-phase extraction cartridges. Desalted peptides were enriched using an Fe-NTA–based immobilized metal affinity chromatography (IMAC) approach (High-Select™ Fe-NTA kit, Thermo Scientific) according to the manufacturer’s instructions. Briefly, peptides were resuspended in binding buffer and incubated with the IMAC resin. After extensive washing, phosphopeptides were eluted under basic conditions. Eluted phosphopeptides were subsequently desalted using peptide desalting spin columns and dried by vacuum centrifugation. For deep proteome coverage, peptides were fractionated using Pierce™ High PH Reversed-Phase Peptide Fractionation Kit (Thermo Scientific). Fractions were concatenated into six pooled fractions, acidified with formic acid to PH < 3, desalted by C18, and dried by vacuum centrifugation prior to LC–MS analysis. Samples were analyzed on an Orbitrap Eclipse Tribrid mass spectrometer coupled to an Easy-nLC 1200 system (Thermo Scientific). For each fraction, ∼2 μg peptides in 1% ACN and 1% FA were loaded onto an in-house packed 100-μm capillary column containing 35 cm Accucore 150 resin (2.6 μm, 150 Å). Peptides were separated over a 180 min gradient from 2% to 23% ACN in 0.125% FA at 500 nL/min. FAIMS Pro (Thermo Scientific) was operated with default settings at compensation voltages of −40, −60, and −80 V. At each voltage, data were acquired in top 10 DDA mode over m/z 400–1400, with MS1 resolution set to 120,000. Singly charged ions were excluded, and multiply charged precursors were fragmented for MS2 using standard automatic gain control and 35% normalized collisional energy (NCE), with dynamic exclusion of 30 s. TMT reporter ions were quantified using multinotch SPS-MS3 (*92*) with 45% NCE.

#### Database searching and quantification

Raw files were converted to mzXML and searched using Comet (*93*) with a previously described in-house pipeline (*94*) against reviewed Mus musculus UniProt entries (downloaded in 2025), supplemented with reversed sequences and common contaminants. Search parameters included 25 ppm precursor mass tolerance, 1.0 Da fragment ion tolerance, full tryptic specificity, and up to two missed cleavages. Methionine oxidation (+15.9949) was set as a variable modification, and TMTpro labeling of lysine residues and peptide N termini (+304.2071) as a static modification. Peptide- and protein-level FDR were controlled below 1%. Phosphorylation site localization was assigned using ModScore with 99% confidence. TMT reporter ions were extracted with a 0.003 Da window, and the most intense peak was used for quantification. Isotopic impurity correction was applied according to the manufacturer’s instructions, and signal-to-noise ratios were calculated. Peptides were excluded if summed S/N was <160 for TMTpro16. NE-responsive sites were identified by comparing NE-treated and control samples at 37°C by two-sided Student’s *t*-test (Benjamini–Hochberg–adjusted *P* < 0.05, fold change > 1). For these sites, fold change between 27°C and 37°C were calculated, with significance assessed by BH-adjusted *t*-tests.

### Histology

Adipose tissues were fixed in 4% paraformaldehyde at 4 °C overnight, followed by incubation in 70% ethanol. After the dehydration procedure, tissues were embedded in paraffin and cut into sections at 5 μm thickness. The sections were stained with haematoxylin and eosin (H&E) according to the standard protocol at the BIDMC pathology core. Images were acquired using the Revolve microscope (ECHO Laboratories) or the Olympus slide scanner.

### Immunostaining

Cells were seeded and differentiated in glass-bottom dishes (VWR, 10810-054) for 4 days. For temperature stimulation, cells were incubated at 27 °C for 90 min, followed by treatment with norepinephrine (NE, 0.1 μM) or PBS for 10 min. Cells were then fixed with 4% paraformaldehyde (PFA) at room temperature for 30 min, washed three times with PBS, and subsequently permeabilized and blocked in PBS containing 0.05% saponin and 5% goat serum for 2 h at room temperature. Samples were incubated with anti–phospho-perilipin 1 (Ser497) antibody in blocking buffer at 4 °C overnight, followed by three washes with PBS. Cells were then incubated with a fluorophore-conjugated goat anti-mouse secondary antibody for 2 h at room temperature and washed three times with PBS. Neutral lipids were stained with BODIPY–FITC for 10 min, followed by nuclear staining with DAPI. Images were acquired using a Zeiss LSM900 confocal microscope and processed using ZEN software (Zeiss). For three-dimensional image analysis, z-stacks were acquired at 0.3-µm intervals and reconstructed using ZEN software. For visualization, lipid-droplet and phospho–perilipin 1 positive regions were manually traced from the corresponding three-dimensional reconstructions, as described previously (*95, 96*).

### Transmission electron microscopy

Tissues were immersion fixed in 2.5% glutaraldehyde (Electron Microscopy Sciences, Hatfield, PA), 2% paraformaldehyde (Electron Microscopy Sciences), 1% osmium tetroxide (Electron Microscopy Sciences) in 0.1M Sodium Cacodylate (Sigma-Aldrich) PH 7.4 on ice for 1 hour. Tissues were grossed immediately to approximately 300um in thickness and less than 2 mm in length and width for high pressure freezing. Tissues were washed with 0.1M Sodium Cacodylate, and then immersed in a 20% fetal bovine serum (Glibco) solution as a cryo-protectant. The tissues were high pressure frozen using a Wohlwend Compact 02 HPF machine (Sennwald) and stored in liquid nitrogen. The tissues were then processed by rapid freeze substitution in acetone containing 1% osmium tetroxide, 0.1% uranyl acetate (Electron Microscopy Sciences) and 5% water for 3 hours before infiltration with LX112 resin (Ladd Research Industries), which was followed by embedding in flat bottom Beem capsules (Electron Microscopy Sciences). The resulting blocks were then sectioned using a Leica Ultracut E ultramicrotome (Leica Microsystems) and sections placed on formvar and carbon-coated grids (Electron Microscopy Sciences). The sections were contrast stained with 2% uranyl acetate followed by lead citrate (Sigma-Aldrich), and imaged in a JEOL JEM 120i transmission electron microscope (JEOL) equipped with a AMT NanoSprint 15-MKII camera (Advanced Microscopy Techniques). Tissue processing, sectioning, staining, and imaging were performed at the BIDMC Electron Microscopy Core.

### BODIPY staining and quantification

Fully differentiated adipocytes were subjected to norepinephrine (NE) or temperature treatments for 6–10 h as indicated. Cells were incubated with BODIPY–FITC (1 μg ml⁻¹, Sigma-Aldrich) at 37 °C for 10 min and immediately imaged using a Revolve fluorescence microscope (ECHO Laboratories). Fluorescence intensity was quantified per field of view using ImageJ software (NIH) and normalized to the corresponding control group for each experiment.

### Immunoblotting

Cells or adipose tissue samples were lysed in lysis buffer (50 mM Tris-HCl, PH 7.4, 150 mM NaCl, 1% NP-40) supplemented with EDTA-free protease inhibitor cocktail (Roche), phosphatase inhibitor cocktails 2 and 3 (Sigma-Aldrich). Lysates were incubated on ice for 10 min and cleared by centrifugation. Protein concentration was determined using a BCA assay (Thermo Fisher Scientific). Equal amounts of protein (20 μg per lane) were mixed with Laemmli sample buffer, denatured by heating, and resolved by SDS–PAGE. Proteins were transferred onto PVDF membranes. Membranes were blocked using EveryBlot Blocking Buffer (Bio-Rad), incubated with primary antibodies at 4 °C overnight, followed by incubation with HRP-conjugated secondary antibodies for 2 h at room temperature. Signals were detected using enhanced chemiluminescence (ECL, Bio-Rad). Vinculin was used as a loading control. Band intensities were quantified using ImageJ software (NIH) and normalized to the corresponding control group.

### RT–qPCR

Total RNA was isolated from cells or tissues using TRIzol reagent (Invitrogen) in combination with the Direct-zol RNA Miniprep kit (R2052, Zymo Research) according to the manufacturer’s instructions. RNA was reverse-transcribed into cDNA using the iScript cDNA Synthesis Kit (Bio-Rad). Quantitative PCR was performed using a QuantStudio 6 system (Applied Biosystems). Gene expression levels were normalized to *β-Actin* as an internal control. Primer sequences used in this study are listed in **Table S3**.

### Statistics

Statistical analyses were performed using a GraphPad Prism v.10 (GraphPad). All data are represented as mean ± s.e.m. unless otherwise specified. Data were obtained from biologically independent samples. Unpaired Student’s *t*-tests were used for two-group comparisons. One-way ANOVA with Tukey’s multiple comparisons test was used for multiple-group comparisons. Two-way ANOVA followed by Šídák’s multiple comparisons test was used to analyze the effects of two independent factors. The statistical parameters and sample numbers are provided in the figure legends. *P* < 0.05 was considered statistically significant unless otherwise specified.

## ACKNOWLEDGMENTS

We are grateful to Hiroshi Nishida, Daisuke Katoh, Beste Noyan, Rosa Li, and Da-qing Wang for their technical support. We also thank Dr. Evan Rosen and Dr. Brad Lowell at Beth Israel Deaconess Medical Center. This work was supported by grants from the National Institutes of Health (NIH) (DK097441 and DK127575) and Howard Hughes Medical Institute to S.K. Y. C. was supported by the American Diabetes Association postdoctoral fellowship (1-26-PDF-0562). Schematic illustrations were created with BioRender.com.

## AUTHORS CONTRIBUTION

Conceptualization, Y.C., S.K.; methodology, Y.C.; investigation, Y.C., L.C., Z.Z., T.L., Y.G., and Z.Y.L.; validation, Y.C. L.C., and Z.Z.; formal analysis, Y.C.; writing – original draft, Y.C. and S.K.; writing – review and editing, all the authors.; funding acquisition, S.K.; supervision, S.K.

## DATA AVAILABILITY

Metabolomics LC–MS raw data have been deposited in the Metabolomics Workbench (https://www.metabolomicsworkbench.org) under project ID ST004745. Phosphoproteomics data are provided in **Table S1**. RNA-seq data have been deposited in the Gene Expression Omnibus under accession number GSE326346. All other data are available in the manuscript or supplementary materials. All the materials and mice are available after completion of MTA.

---

**Supplementary Figure 1.**
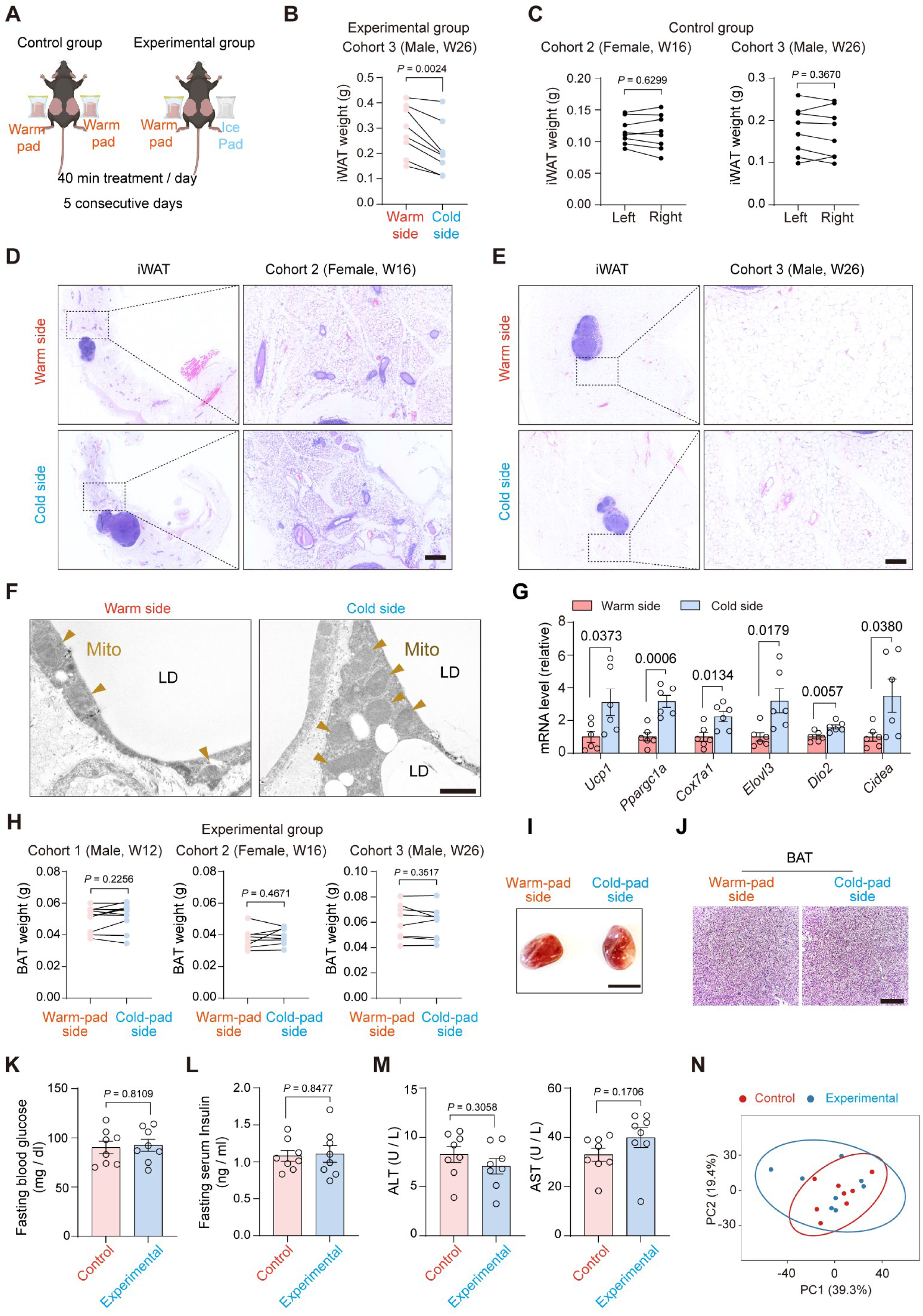
The validation of tissue cooling effect on adipose tissue mass and systemic metabolic parameters. **A.** Schematic illustration of the local adipose tissue cooling experiments. Mice housed at room temperature were shaved on both inguinal sides. Under anesthesia, the control group (<u>Left</u>) was placed on warm pads on both inguinal sides, whereas the experimental group (<u>Right</u>) was placed on a warm pad on one inguinal side and an ice pad on the other side for 10 min. This procedure was repeated 4 times per day, resulting in a total of 40 min of local cold exposure per day for 5 consecutive days. **B.** iWAT weight of Cohort 3 (26-week-old male mice) following 5-day treatment with ice/warm pad. Each dot represents one depot, with matched warm- and cold-treated depots from the same mouse connected by lines. *n* = 8. Paired *t*-test. **C.** iWAT weight of the control group mice that received warm pads on both inguinal sides. Each dot represents one depot, with matched depots from the same mouse connected by lines. <u>Left,</u> cohort 2 (16-week-old female mice); <u>Right</u>, cohort 3 (26-week-old male mice). *n* = 8 mice per cohort. Paired *t*-test. **D.** Representative H&E staining of iWAT sections from cohort 2 that received either a warm pad or an ice pad. <u>Left</u>, whole-slide scans; <u>Right</u>, magnified views of the indicated regions. Scale bar: 200 μm. **E.** Representative H&E staining of iWAT sections from cohort 3 that received either a warm pad or an ice pad. <u>Left</u>, whole-slide scans; <u>Right</u>, magnified views of the indicated regions. Scale bar: 200 μm. **F.** Representative transmission electron microscopy images of iWAT that received either a warm pad or an ice pad. Arrowheads indicate mitochondria. Scale bar: 1 μm. **G.** mRNA levels of thermogenic genes in iWAT from mice that received local cold-pad or warm-pad treatment for 5 days. Gene-expression levels were normalized to the warm-pad control group. *n* = 6 mice per group. Data are mean ± s.e.m. Unpaired *t*-test. **H.** Interscapular BAT weight of three independent cohorts that received warm pads on both inguinal sides. Each dot represents one BAT lobe, with matched warm- and cold-treated depots from the same mouse connected by lines. Cohort 1, 12-week-old male mice (*n* = 11); Cohort 2, 16-week-old female mice (*n* = 8); Cohort 3, 26-week-old male mice (*n* = 8). Paired *t*-test. **I.** Representative tissue images of interscapular BAT depots from mice in (**H**). Scale bar, 0.5 cm. **J.** Representative hematoxylin and eosin (H&E) staining of BAT sections in (**H**). Scale bar, 210 μm. **K.** Fasting blood glucose levels in cohort 3 mice. The control group (both warm pads) and the experimental group (one ice pad and one warm pad) were fasted for 6 h before measurement. *n* = 8. Unpaired *t*-test. **L.** Fasting serum insulin levels in cohort 3 mice. The control and experimental groups were fasted for 6 h before measurement. *n* = 8. Unpaired *t*-*t*est. **M.** Serum alanine aminotransferase (ALT; left) and aspartate aminotransferase (AST; right) levels in cohort 2 mice. *n* = 8. Unpaired *t*-test. **N.** Principal component analysis (PCA) of global serum lipid species in cohort 2 mice. *n* = 8.

**Supplementary Figure 2.**
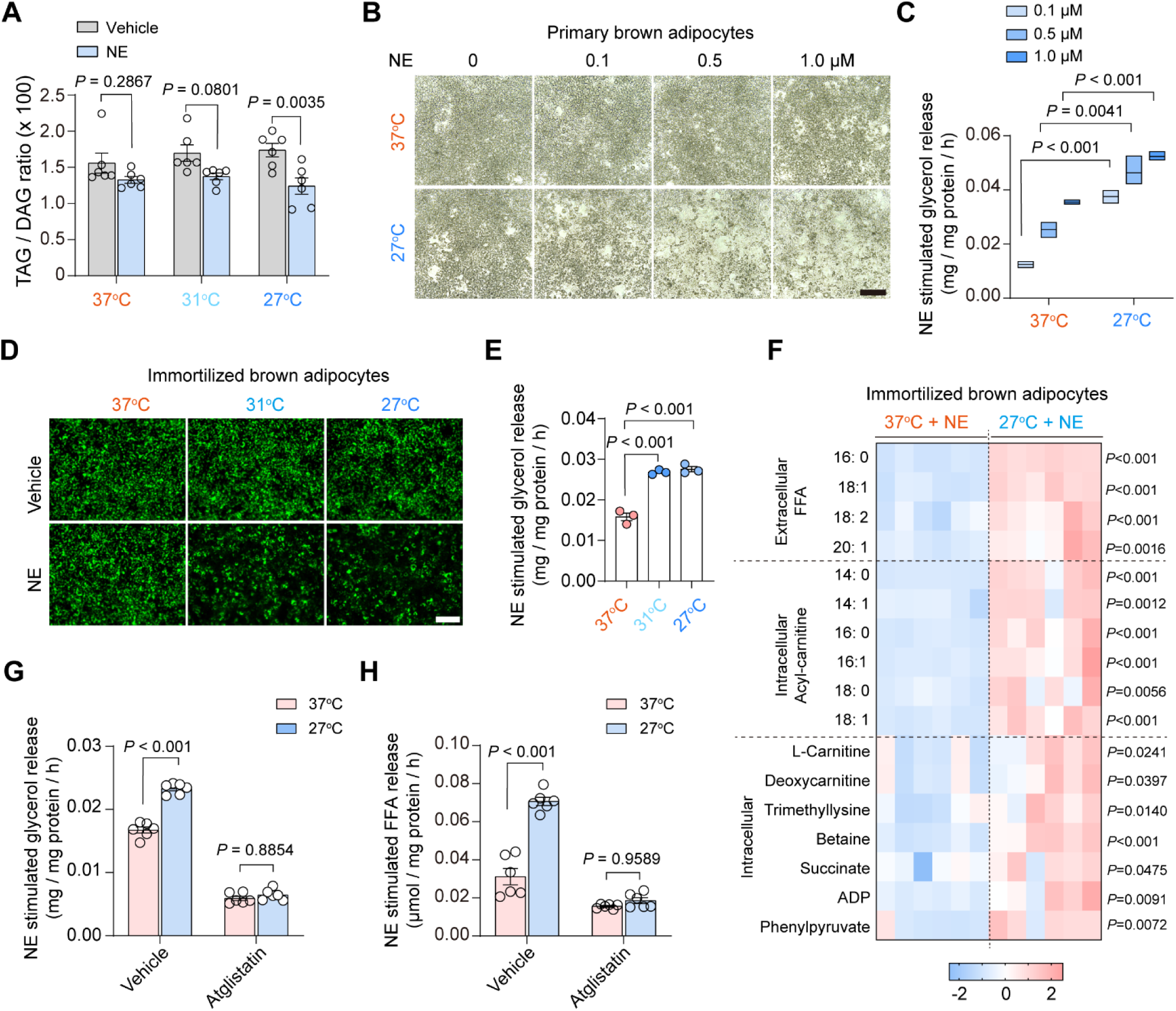
Cell-autonomous enhancement of lipolysis by cooling temperatures. **A.** The ratio of cellular triacylglycerol to diacylglycerol (DAG) in primary inguinal adipocytes treated at the indicated temperatures (37 °C, 31 °C, and 27 °C) in the presence of vehicle or norepinephrine (NE) at 0.1 μM. *n* = 6. Data are presented as mean ± s.e.m. Two-way ANOVA with Šídák’s multiple comparisons test. **B.** Representative bright-field images of differentiated primary brown adipocytes treated with increasing concentrations of norepinephrine (NE) at 0.1, 0.5, and 1.0 μM at 37 °C or 27 °C for 10 hours. Scale bar: 210 µm. **C.** Glycerol release from differentiated primary brown adipocytes treated with increasing concentrations of NE (0.1, 0.5, and 1.0 μM) at 37 °C or 27 °C. Lipolysis was quantified as glycerol release normalized to protein amount. *n* = 3. Data are presented as mean ± s.e.m. Unpaired *t*-test. **D.** Representative BODIPY staining of differentiated brown adipocytes treated at the indicated temperatures (37 °C, 31 °C, and 27 °C) with vehicle or norepinephrine (NE) at 0.1 μM. Scale bar: 130 µm. **E.** Glycerol release from differentiated brown adipocytes treated at the indicated temperatures (37 °C, 31 °C, and 27 °C) with vehicle or norepinephrine (NE) at 0.1 μM. Lipolysis was quantified as glycerol release normalized to total protein. *n* = 3. Data are presented as mean ± s.e.m. One-way ANOVA followed by Tukey’s multiple comparisons test. **F.** Heatmap showing changes in medium BSA-bound free fatty acids (FFA), cellular acyl-carnitines and cellular indicated metabolites following NE treatment for 10 hours at 27°C or 37°C. Values are presented as Z-scores. *n* = 6 per group. Unpaired *t*-test. **G.** Glycerol release from differentiated brown adipocytes treated at 37 °C or 27 °C with vehicle (DMSO) or Atglistatin (20 μM), in the presence or absence of norepinephrine (NE) at 0.1 μM for 10 h. *n* =6 per group. Data are normalized to total protein and presented as mean ± s.e.m. Two-way ANOVA with Šídák’s multiple comparisons test. **H.** Free fatty acids (FFA) release from immortalized brown adipocytes treated at 37 °C or 27 °C with vehicle (DMSO) or Atglistatin (20 μM), in the presence or absence of norepinephrine (NE, 0.1 μM) for 10 h. *n* =6 per group. Data are normalized to total protein and presented as mean ± s.e.m. Two-way ANOVA with Šídák’s multiple comparisons test.

**Supplementary Figure 3.**
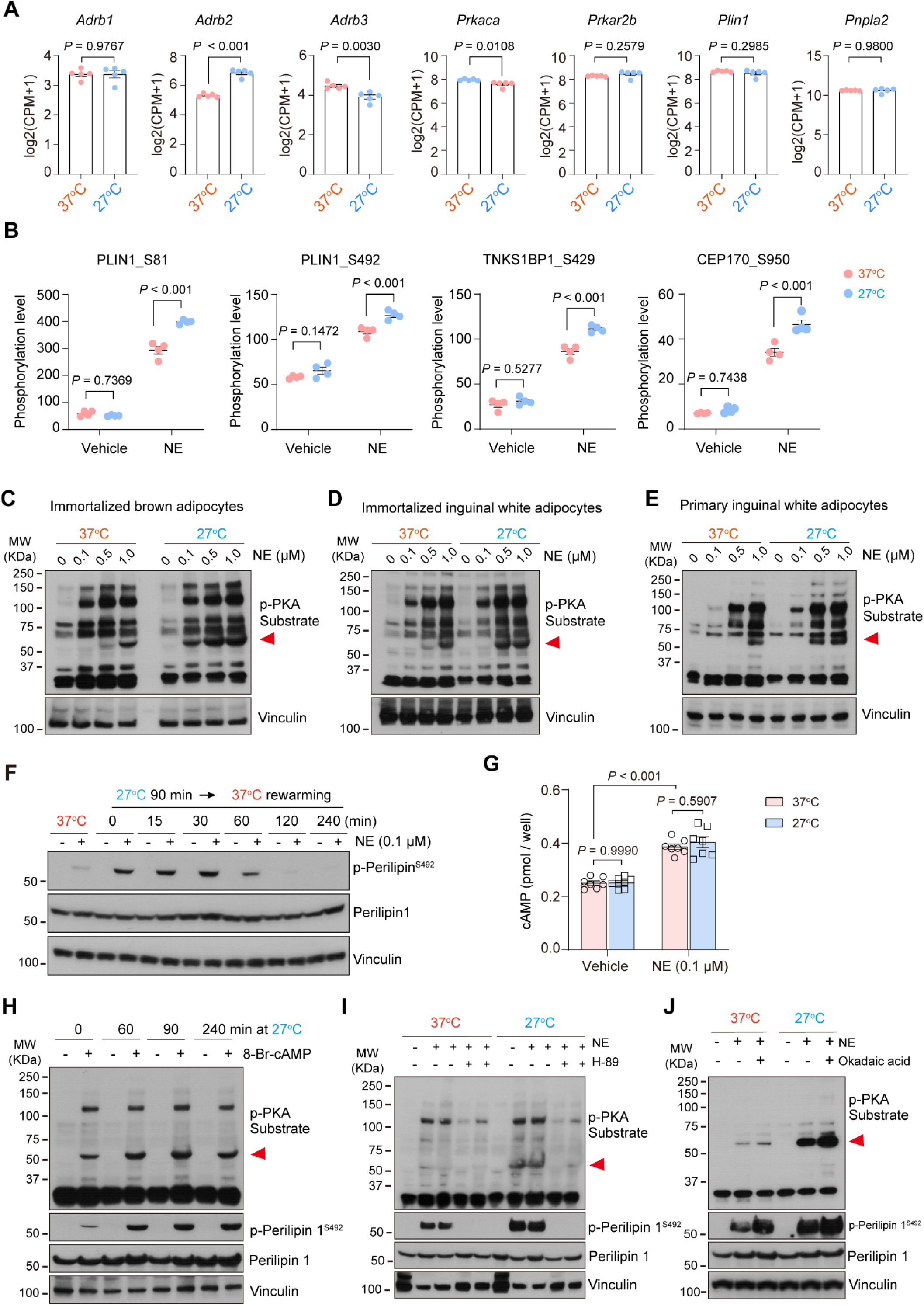
Cooling temperatures selectively amplify PKA-perilipin 1 signaling. **A.** Immortalized brown adipocytes were pre-incubated at 27 °C for 1 h prior to stimulation with norepinephrine (NE) at 0.1 μM for 3 h. Cells were harvested for RNA sequencing (Plasmidsaurus). Expression levels of genes related to lipolysis are presented as log2-transformed CPM values log2(CPM + 1). *n* = 5. Data are shown as mean ± s.e.m.. Statistical significance was determined using an unpaired two-tailed Student’s *t*-test on log-transformed CPM values. **B.** Quantification of phosphorylation levels of representative sites identified from phosphoproteomic analysis, including perilipin 1 (S81 and S492), TNKS1BP1 (S429), and CEP170 (S950), under vehicle or norepinephrine (NE) at 0.1 μM stimulation at 37 °C or 27 °C. *n* = 4 per group. Data are presented as mean ± s.e.m. Two-way ANOVA with Šídák’s multiple comparisons test. **C.** Differentiated brown adipocytes were pre-incubated at 37 °C or 27 °C for 90 min, followed by stimulation with norepinephrine (NE) at 0.1, 0.5, and 1.0 μM for 10 min. Immunoblot analysis shows phosphorylation of the indicated PKA substrates. Vinculin was used as a loading control. **D.** Immortalized inguinal WAT-derived differentiated adipocytes were treated as in (**C**). Immunoblot analysis shows phosphorylation of PKA substrates. Vinculin was used as a loading control. **E.** Primary inguinal WAT-derived adipocytes were treated as in (**C**). Immunoblot analysis shows phosphorylation of PKA substrates. Vinculin was used as a loading control. **F.** Reversibility of cooling-induced perilipin 1 phosphorylation after rewarming. Immunoblot analysis of perilipin 1^S492^ phosphorylation in adipocytes following 90 min cooling at 27°C and subsequent rewarming at 37°C for up to 240 min. Vinculin was used as a loading control. **G.** Intracellular cAMP levels in adipocytes incubated at 27°C or 37°C for 90 min. Cells were treated with norepinephrine (NE; 0.1 μM) for 10 min before harvest. No IBMX is used to inhibit phosphodiesterase activity. *n* = 7 per group. Data are presented as mean ± s.e.m. Two-way ANOVA with Šídák’s multiple comparisons test. **H.** Differentiated brown adipocytes were pre-incubated at 27 °C for 0 min, 60 min, 90 min or 240 min, followed by stimulation with 8-Br-cAMP (200 μM) for 10 min. Immunoblot analysis shows phosphorylation of PKA substrates, phospho-perilipin 1^S492,^ and total perilipin 1. Vinculin was used as a loading control. **I.** Differentiated brown adipocytes were pre-incubated at 37 °C or 27 °C for 90 min and then treated with the PKA inhibitor H-89 (15 μM) or vehicle for 20 min before stimulation with norepinephrine (NE; 0.1 μM) or vehicle. Immunoblot analysis shows phosphorylation of PKA substrates, phospho-perilipin 1^S492,^ and total perilipin 1. Vinculin was used as a loading control. **J.** Differentiated brown adipocytes were pre-incubated at 37 °C or 27 °C for 60 min, followed by treatment with okadaic acid (50 nM) for 30 min. Cells were then stimulated with norepinephrine (NE) at 0.1 μM for 10 min prior to harvest. Immunoblot analysis shows phosphorylation of PKA substrates, phospho-perilipin 1^S492,^ and total perilipin 1. Vinculin was used as a loading control.

**Supplementary Figure 4.**
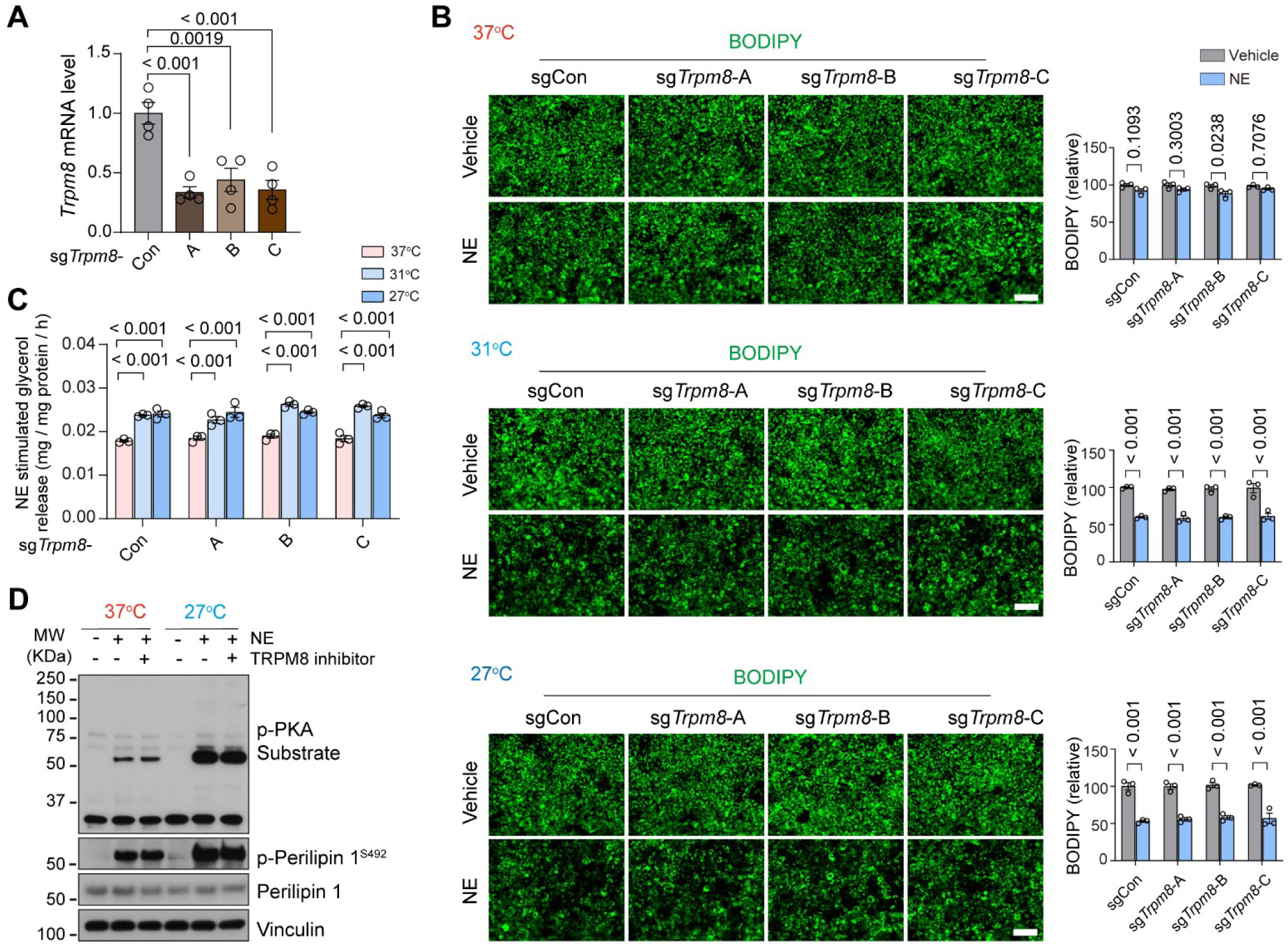
Cooling temperatures selectively augment PKA signaling independent of the TRP channels. **A.** mRNA levels of *Trpm8* in adipocytes expressing control sgRNA (sgCon) or three independent sgRNAs targeting *Trpm8* (sg*Trpm8*-A, -B, and -C). *n* = 4 per group. Data are presented as mean ± s.e.m. One-way ANOVA followed by Tukey’s multiple comparisons test. **B.** Representative BODIPY staining of adipocytes expressing sgCon or sg*Trpm8* constructs under vehicle or norepinephrine (NE) at 0.1 μM for 10 hours at the indicated temperatures. Scale bar: 130 µm. Quantification of BODIPY fluorescence intensity is shown on the right. *n* = 3 per group. Data are presented as mean ± s.e.m. Two-way ANOVA with Šídák’s multiple comparisons test. **C.** NE-induced glycerol release in adipocytes expressing sgCon or sg*Trpm8* constructs treated at 37 °C, 31 °C, or 27 °C after NE treatment at 0.1 μM for 10 hours. *n* = 3 per group. Data are presented as mean ± s.e.m. Two-way ANOVA with Šídák’s multiple comparisons test. **D.** Differentiated brown adipocytes were pre-treated with the TRPM8 inhibitor BCTC (10 μM) or vehicle (DMSO) for 10 min, followed by incubation at 37 °C or 27 °C for 90 min and subsequent stimulation by norepinephrine (NE) at 0.1 μM for 10 min. Immunoblot analysis shows phosphorylation of PKA substrates, phospho-perilipin 1^S492,^ and total perilipin 1. Vinculin was used as a loading control.

**Supplementary Figure 5.**
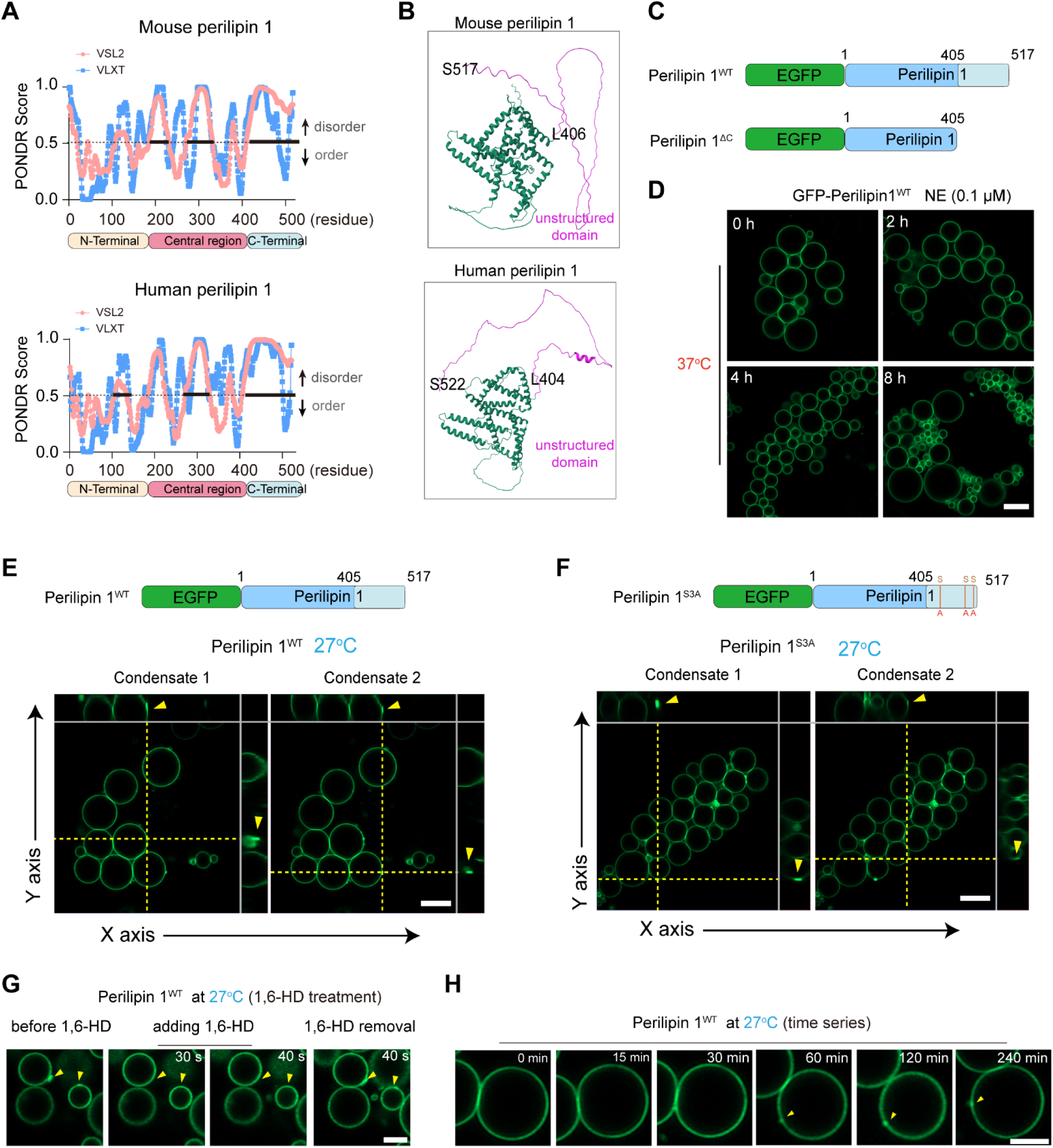
Cooling temperatures induce the formation of perilipin 1 condensates. **A.** Prediction of intrinsically disordered regions in mouse (<u>Top</u>) and human (<u>Bottom</u>) perilipin 1 using the PONDR algorithm (pink: VSL2, blue: VLXT) with bold lines indicating predicted disordered regions. The bottom schematic illustrates domain annotations of the human perilipin 1 protein (N-terminal, Central region, and C-terminal). **B.** Structural prediction of the mouse (<u>Top</u>) and human (<u>Bottom</u>) perilipin 1 protein by Alphafold3, highlighting a folded N-terminal domain and an extended C-terminal unstructured region. Residues after L406 (mouse) or L404 (human) form a predicted intrinsically disordered region (IDR), lacking defined secondary structure. **C.** Schematic of GFP-tagged perilipin 1 constructs, including full-length perilipin 1 (Perilipin 1^WT^), C-terminal truncation (Perilipin 1^ΔC^). **D.** Representative GFP-perilipin 1 images in differentiated adipocytes treated with norepinephrine (NE; 0.1 μM) at 37°C for the indicated times, up to 8 h. Scale bar: 4 μm. **E.** <u>Top</u>: Schematic of GFP-tagged full-length perilipin 1 (Perilipin 1^WT^). <u>Bottom</u>: Representative orthogonal visualizations of indicated condensates at 27°C, reconstructed from confocal Z-stack images. Yellow arrows indicate perilipin1 condensates. Scale bar: 4 μm. **F.** <u>Top</u>: Schematic of GFP-tagged C-terminal PKA phosphorylation-deficient mutant (Perilipin 1^S3A^; S433A/S492A/S517A). <u>Bottom</u>: Representative orthogonal visualizations of indicated condensates at 27°C, reconstructed from confocal Z-stack images. Yellow arrows indicate perilipin1 condensates. Scale bar: 4 μm. **G.** Cooling-induced perilipin 1 puncta formation in response to 1,6-hexanediol (1,6-HD, 5%) treatment. Representative images showing GFP-perilipin 1 puncta in adipocytes treated at 27 °C before, after 1,6-HD treatment for 30 or 40 seconds, and after the withdrawal of 1,6-HD. Yellow arrows indicate perilipin1 condensates. Scale bar: 2 μm. **H.** Time-lapse live-cell imaging of GFP-perilipin 1 condensate formation at 27°C. Time points are shown on the right. Yellow arrows indicate perilipin1 condensates. Scale bar: 2 μm.

**Supplementary Figure 6.**
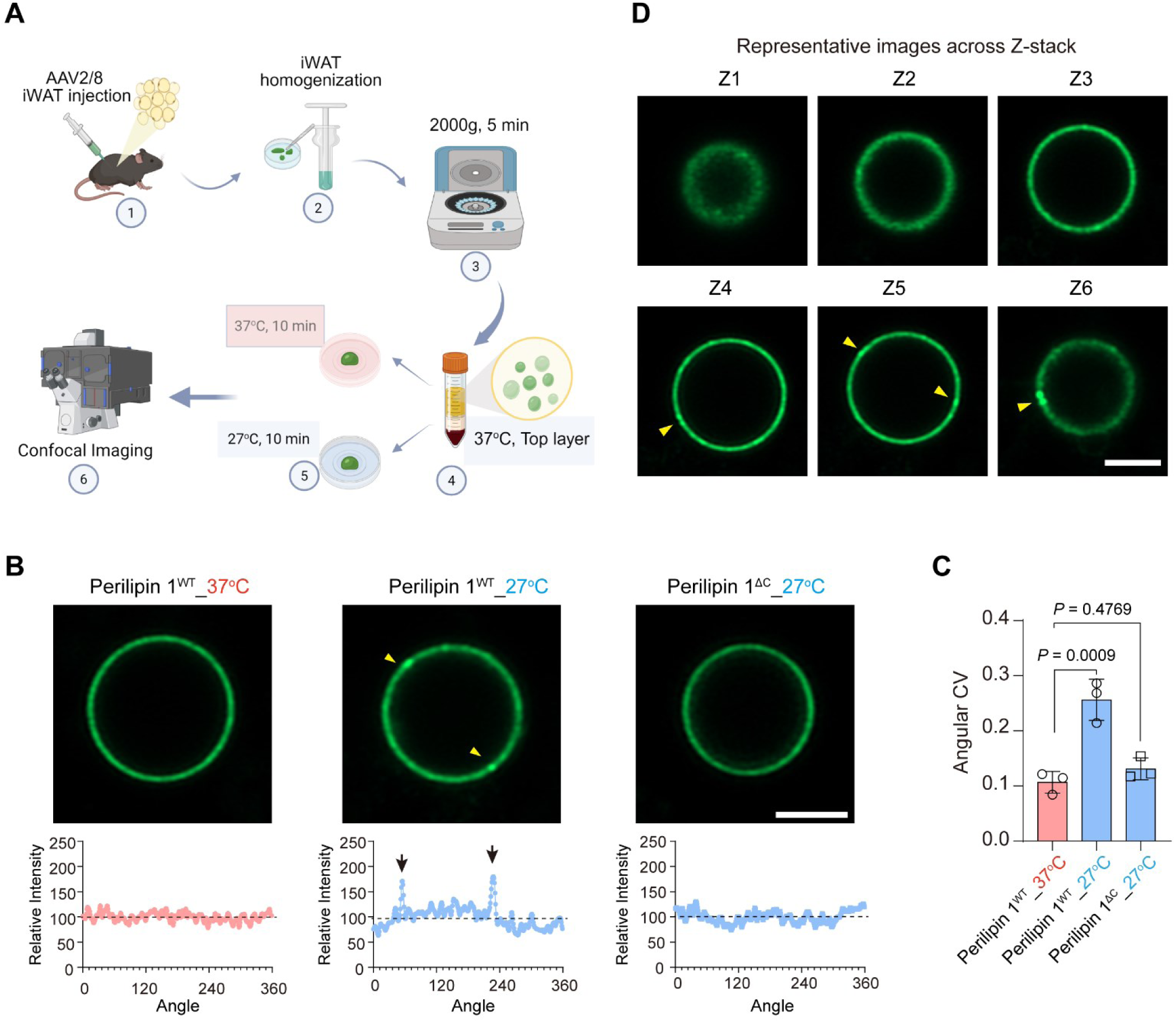
Cooling induces the C-terminal-dependent perilipin 1 condensate formation on lipid droplets. **A.** Workflow for assessing perilipin 1 condensation on lipid droplets. iWAT of 8-week-old male mice was subcutaneously injected with AAV2/8 expressing GFP-PLIN1. Three weeks after injection, lipid droplets were isolated from iWAT, placed in glass-bottom dishes, and incubated at 37 °C or 27 °C for 10 min, followed immediately by confocal imaging. **B.** <u>Top</u>: Representative GFP images of full-length GFP-PLIN1 (Perilipin 1^WT^) on isolated lipid droplets from cultured adipocytes incubated at 37 °C (left) or 27 °C (middle), and GFP–perilipin 1^ΔC^ on isolated lipid droplets incubated at 27 °C (right). <u>Bottom</u>: Corresponding angular distributions of GFP fluorescence intensity around the lipid droplets, using Fiji software. Arrowheads indicate perilipin 1 condensates on lipid droplets. **C.** Coefficient of variation (CV) of perilipin 1 fluorescence intensity on lipid droplet surface under the indicated conditions. *n*= 3 independent experiment per group. One-way ANOVA followed by Tukey’s multiple comparisons test. **D.** Representative Z-stack images of perilipin 1 condensates from (**B**). Scale bar: 2 μm. Arrowheads indicate perilipin 1 condensates on lipid droplets.

**Supplementary Figure 7.**
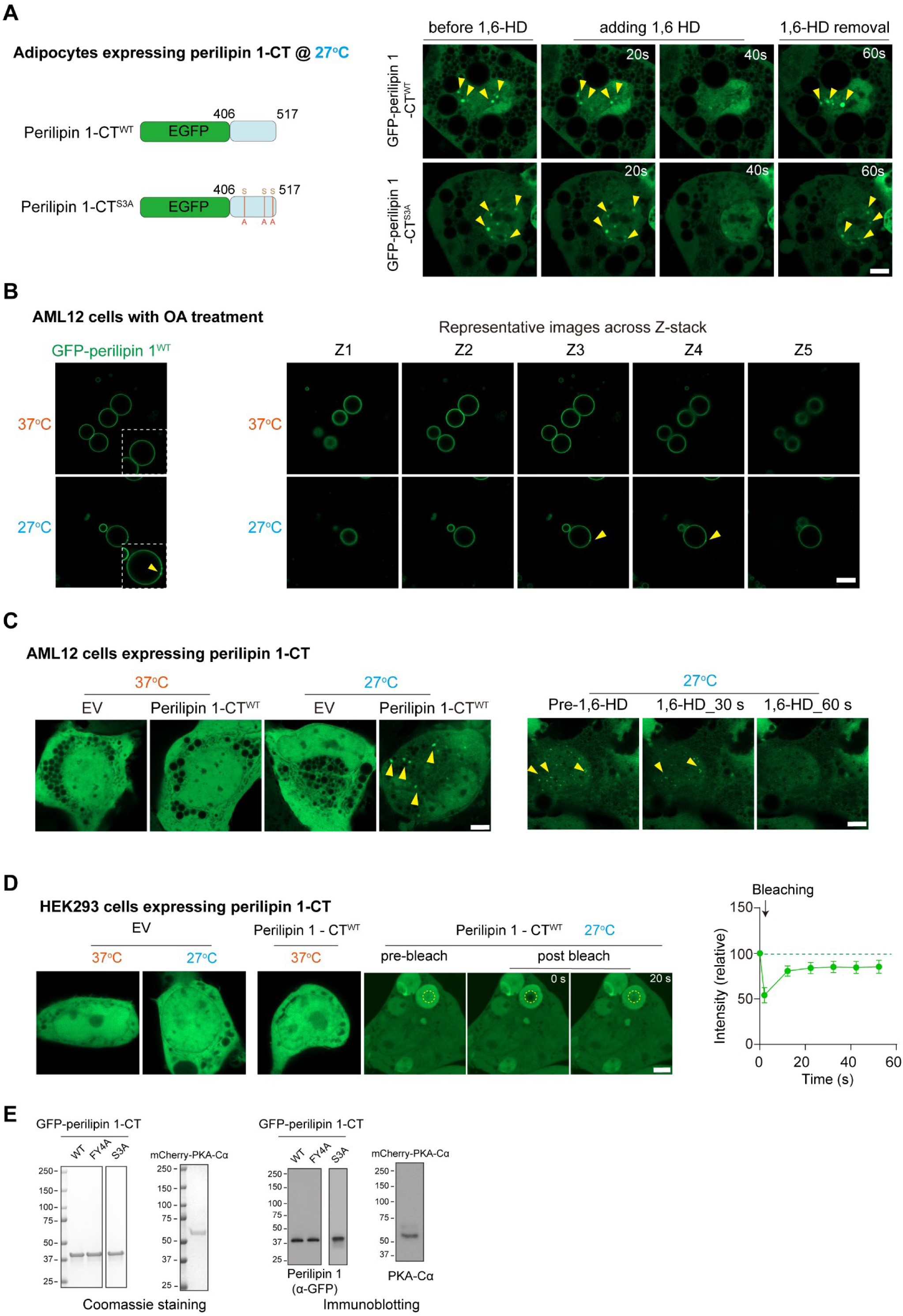
The C-terminal domain of perilipin 1 is sufficient for cooling-induced LLPS. **A.** <u>Left</u>: Schematic of GFP-tagged constructs containing the C-terminal region of perilipin 1. <u>Right</u>: Representative images of adipocytes expressing GFP-perilipin 1-CT^WT^ (Top) or GFP-perilipin 1-CT^S3A^ (S433/492/517A; Bottom) at 27 °C for 8 hours before, during, and after treatment with 1,6-hexanediol (1,6-HD, 5%). Perilipin 1 condensates (arrowheads) rapidly dissolved upon 1,6-HD treatment and reformed after removal. Scale bar: 4 µm. **B.** <u>Left</u>: Representative GFP–perilipin 1 images of AML12 hepatocytes expressing GFP–perilipin 1. Cells were treated with oleic acid (OA) for 48 h to induce lipid-droplet formation and subsequently incubated at 37°C or 27°C. Arrowheads indicate perilipin 1 condensates. <u>Right</u>: Representative serial Z-stack images of corresponding GFP–perilipin 1 hepatocytes. Scale bar: 4 μm. **C.** <u>Left</u>: Representative images of AML12 cells expressing GFP-perilipin 1-CT^WT^ or GFP control after incubation at 37 °C or 27 °C for 8 hours. <u>Right</u>: Representative images of AML12 cells expressing GFP-perilipin 1-CT^WT^ cultured at 27 °C before and after treatment with 1,6-hexanediol (1,6-HD, 5%). Scale bar: 4 μm. Arrowheads indicate perilipin 1 condensates. **D.** <u>Left</u>: HEK293 cells expressing GFP–perilipin 1-CT or GFP control were cultured at 27 °C for 8 h or 37 °C as a control. FRAP analysis of perilipin 1-CT condensates at 27 °C shows rapid fluorescence recovery. Scale bar: 4 µm. <u>Right</u>: Quantification of the bleaching area is shown on the right. *n* = 3. **E.** <u>Left</u>: Coomassie Brilliant Blue staining of purified GFP–perilipin 1-CT^WT^, GFP–perilipin 1-CT^FY4A^, GFP–perilipin 1-CT^S3A^, and mCherry-PKA-Cα. <u>Right</u>: Immunoblotting with anti-GFP or PKA-Cα confirmed protein identity.

**Supplementary Figure 8.**
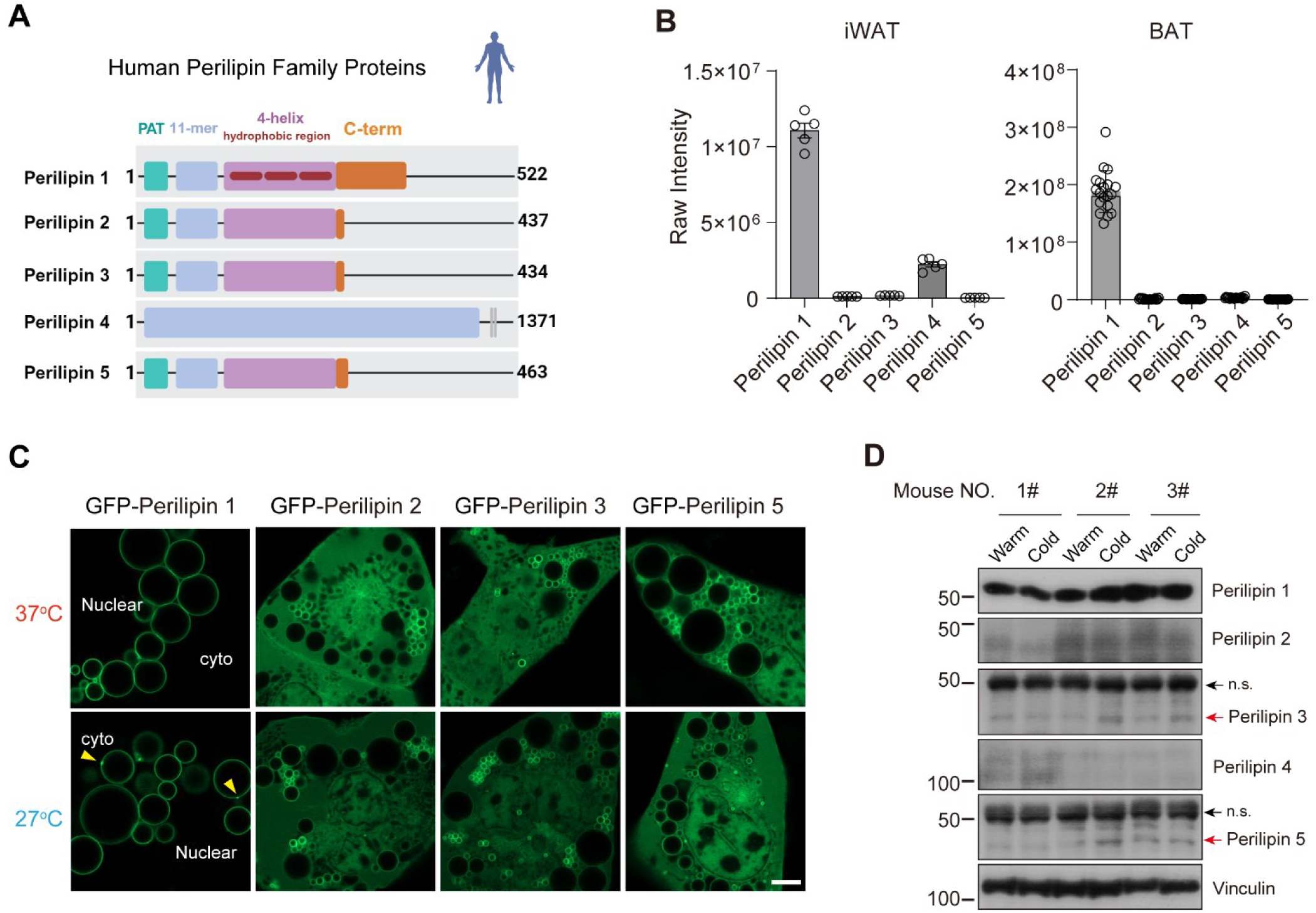
The distinct feature of perilipin 1 among perilipin family proteins. **A.** Schematic structural comparison of human perilipin family members (Perilipin 1, 2, 3, 4, and 5), highlighting the conserved domains, including the PAT domain, 11-mer repeats, and amphipathic helices. Note that perilipin 1 is distinct from other family members in the C-terminal extension and three hydrophobic regions. PLIN4 exhibits an extended central repeat region. Protein lengths are indicated on the right. **B.** <u>Left</u>: Protein abundance (raw intensity) of perilipin family members in iWAT from a publicly available dataset. <u>Right</u>: Protein abundance (raw intensity) of perilipin family members in BAT from a publicly available dataset. **C.** Representative images of adipocytes expressing GFP-tagged perilipin 1, perilipin 2, perilipin 3, or perilipin 5 following incubation at 37°C or 27°C. Arrowheads indicate perilipin 1 condensates on lipid droplets. Scale bar: 4 μm. **D.** Immunoblot analysis showing no noticeable changes in perilipin 1–5 protein expression in iWAT from the ice-pad-treated side (cold) compared with warm-pad-treated side (warm). n.s., non-specific band.

**Supplementary Figure 9.**
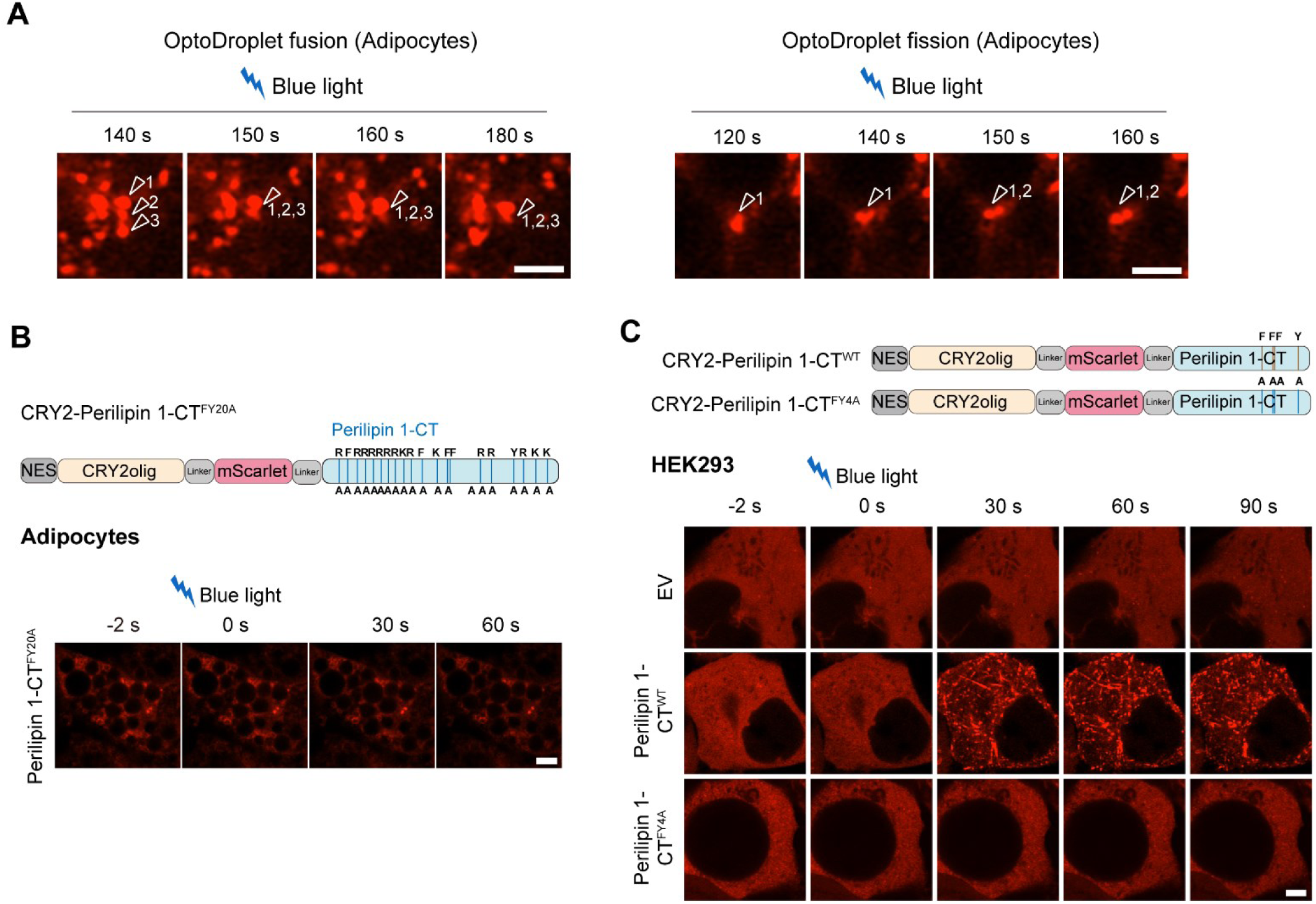
The structural requirement for light-induced perilipin 1 phase separation. **A.** Time-lapse imaging of optogenetic perilipin 1-CT droplets in adipocytes following blue light stimulation (see Figure 4D). <u>Left</u>: Representative images showing fusion of multiple condensates over time from 140 to 180 sec. <u>Right</u>: Representative images showing fission events from 120 to 160 s, where individual condensates split into smaller droplets. Arrowheads indicate tracked condensates. Scale bar: 2 µm. **B.** <u>Top</u>: Schematic of the CRY2-based optogenetic constructs used to test the clustering of a phase separation-deficient perilipin 1 mutant. The mutant (perilipin 1-CT^FY20A^), in which five aromatic residues (F418, F479, F495, F496, Y510) and fifteen lysine/arginine residues (R408, R419, R436, R445, R450, R454, R457, K470, R476, K486, R497, R507, R514, K515, K516) were substituted with alanine, was fused to CRY2olig–mScarlet with a nuclear export signal. <u>Bottom</u>: Representative images in differentiated brown adipocytes expressing perilipin 1-CT^FY20A^ before (−2 s) and after blue light stimulation at 0, 30, and 60 s. Scale bar: 4 µm. **C.** <u>Top</u>: Schematic of the CRY2-based optogenetic constructs used in HEK293 cells. <u>Bottom</u>: Representative images in HEK293 cells expressing empty vector (EV), perilipin 1-CT^WT^, or perilipin 1-CT^FY4A^ before (−2 s) and after blue light stimulation at 0, 30, 60, and 90 s. Scale bar: 4 µm.

**Supplementary Figure 10.**
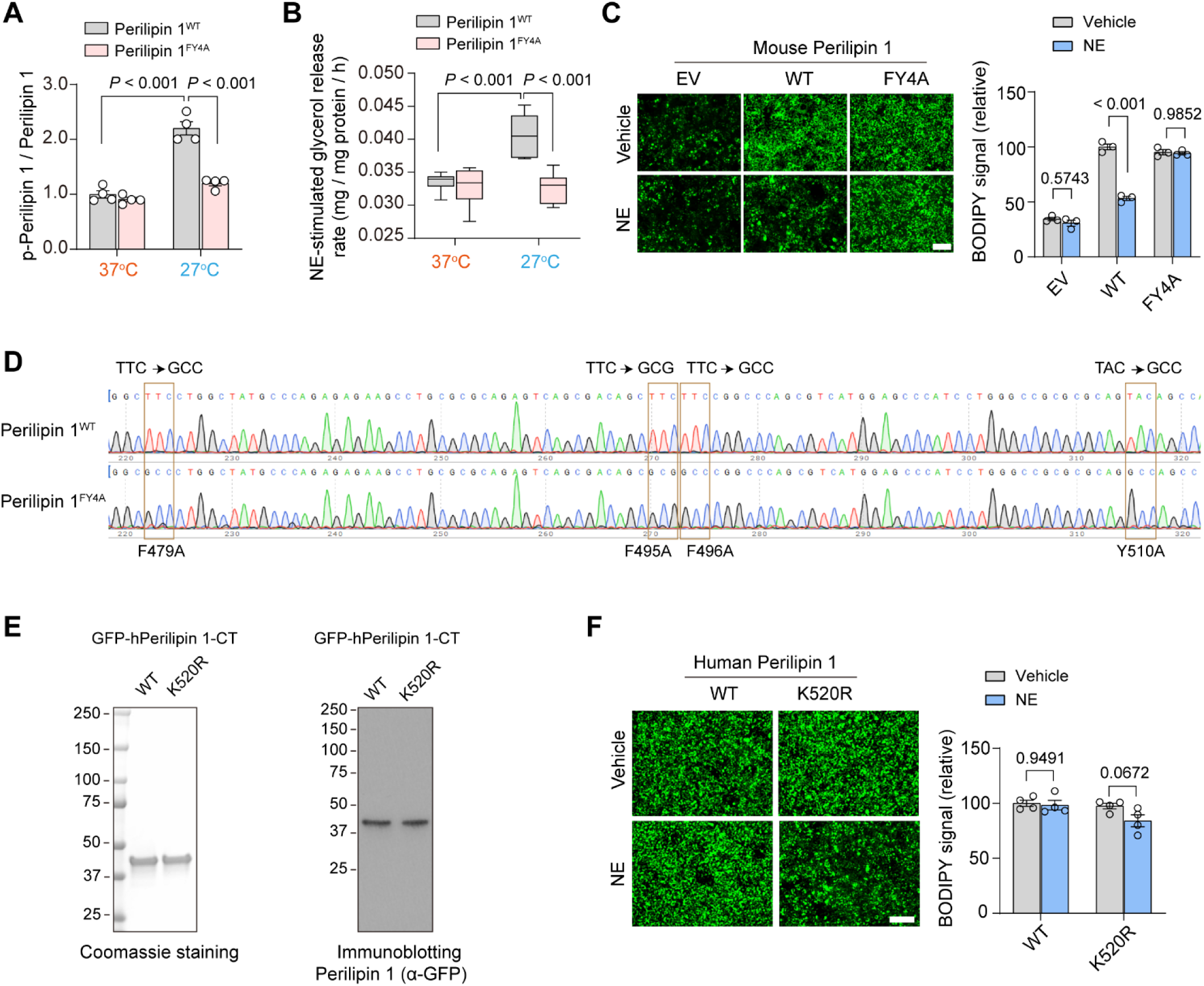
Perilipin 1 LLPS is required for cooling-induced lipolysis. **A.** Quantification of phospho-perilipin 1 levels shown in Fig. 4E. *n* = 4 per group. Data are presented as mean ± s.e.m. Two-way ANOVA with Šídák’s multiple comparisons test. **B.** Lipolysis was assessed by measuring glycerol release into the conditioned media after 8 hours of norepinephrine (NE, 0.1 μM) stimulation. Glycerol levels were normalized to total protein. *n* = 6 per group. Data are presented as mean ± s.e.m. Two-way ANOVA with Šídák’s multiple comparisons test. **C.** <u>Left</u>: Representative BODIPY staining of *Plin1* KO adipocytes reconstituted with an empty vector (EV), mouse perilipin 1^WT^, or the phase separation–deficient mutant perilipin 1^FY4A^ treated with vehicle or norepinephrine (NE, 0.1 μM) at 27 °C. Scale bar: 130 µm. <u>Right</u>: Quantification of BODIPY fluorescence intensity is shown on the right. *n* = 3 per group. Data are presented as mean ± s.e.m. Two-way ANOVA followed by with Šídák’s multiple comparisons test. **D.** Sanger sequencing confirmation of the perilipin 1^FY4A^ knock-in mouse allele, carrying the F479A, F495A, F496A, and Y510A substitutions. **E.** <u>Left</u>: Coomassie Brilliant Blue staining of purified GFP-tagged human perilipin 1 C-terminal proteins (GFP-perilipin 1-CT^WT^ and GFP-perilipin 1-CT^K520R^). <u>Right</u>: Immunoblotting with anti-GFP confirmed their protein identity. **F.** <u>Left</u>: Representative BODIPY staining of *Plin1* KO adipocytes reconstituted with human perilipin 1^WT^ or perilipin 1^K520R^ treated with vehicle or norepinephrine (NE) at 37 °C. Scale bar: 130 µm. <u>Right</u>: Quantification of BODIPY fluorescence intensity. *n* = 4 per group. Data are presented as mean ± s.e.m. Two-way ANOVA followed by Šídák’s multiple comparisons test.

**Supplementary Figure 11.**
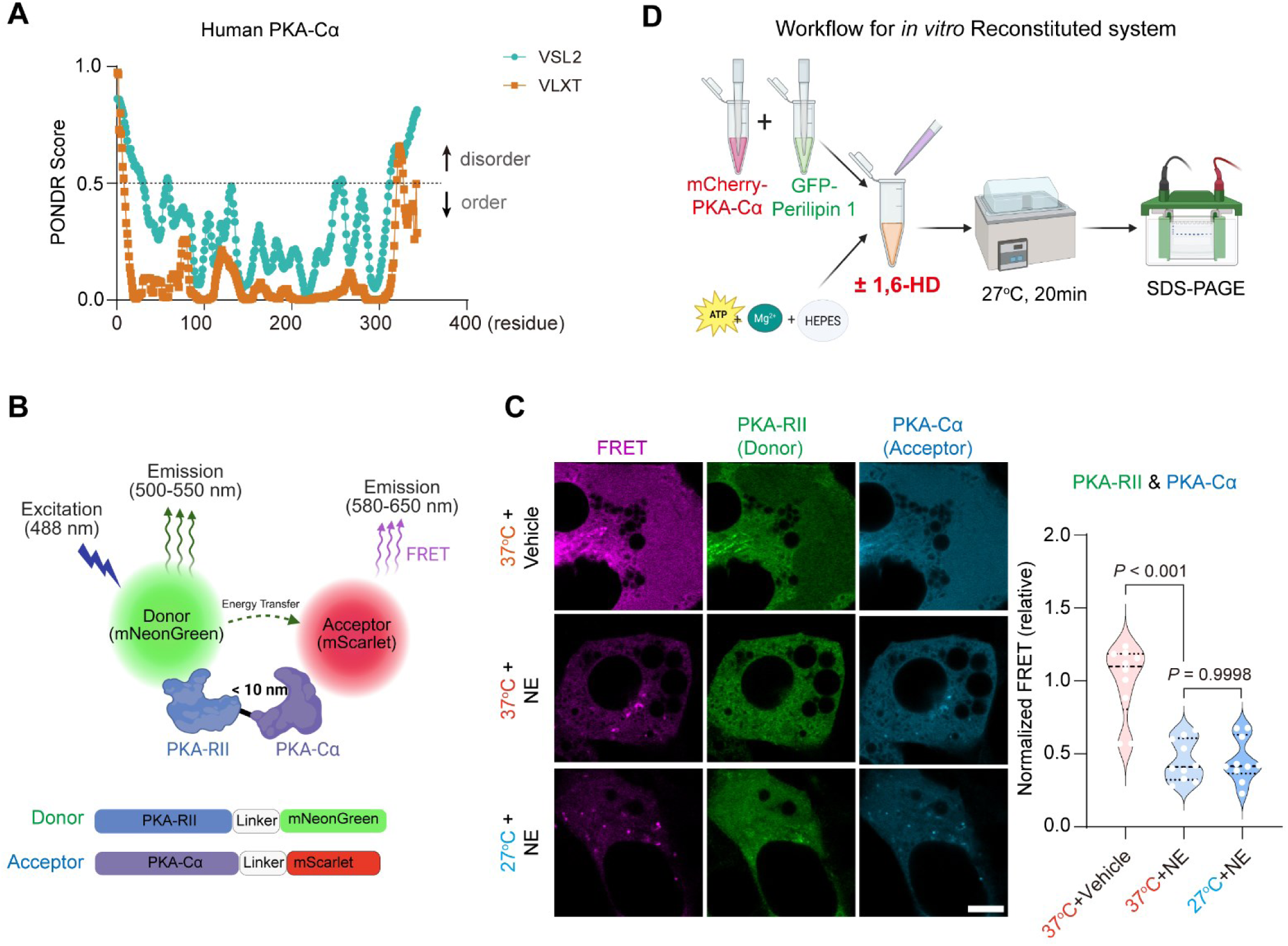
Cooling temperature did not alter the proximity of PKA-Cα and PKA-RII in adipocytes. **A.** Prediction of intrinsically disordered regions in human PKA-Cα using the PONDR algorithm (pink: VSL2, blue: VLXT). **B.** Schematic illustration of the Förster resonance energy transfer (FRET) system using PKA-RII-mNeonGreen as the donor and PKA-Cα–mScarlet as the acceptor. Upon excitation at 488 nm, energy transfer occurs when the two proteins are within <10 nm, leading to acceptor emission (580–650 nm). Donor emission (500–550 nm) and acceptor signals were used to quantify FRET efficiency. **C.** <u>Left</u>: Representative images of FRET-based proximity assay between PKA-RII and PKA-Cα in adipocytes. Differentiated brown adipocytes stably expressing PKA-RII-mNeonGreen (Donor) and PKA-Cα–mScarlet (Acceptor) were incubated at 37 °C or 27 °C for 90 min, followed by NE stimulation at 0.1 μM for 1 min. Scale bar: 5 µm. <u>Right</u>: Quantification of FRET signals. *n* = 10 adipocytes per group. Data are presented as mean ± s.e.m. One-way ANOVA followed by Tukey’s multiple comparisons test. **D.** Workflow of the *in vitro* reconstituted system to assess perilipin 1 phosphorylation by PKA-Cα. Recombinant GFP–perilipin 1 and mCherry–PKA-Cα proteins were mixed in a buffer containing HEPES, Mg²⁺, and ATP, in the presence or absence of 1,6-HD at 5%. Samples were incubated at 27 °C for 20 min prior to SDS–PAGE analysis to assess perilipin 1 phosphorylation under condensate-forming conditions.

**Supplementary Figure 12.**
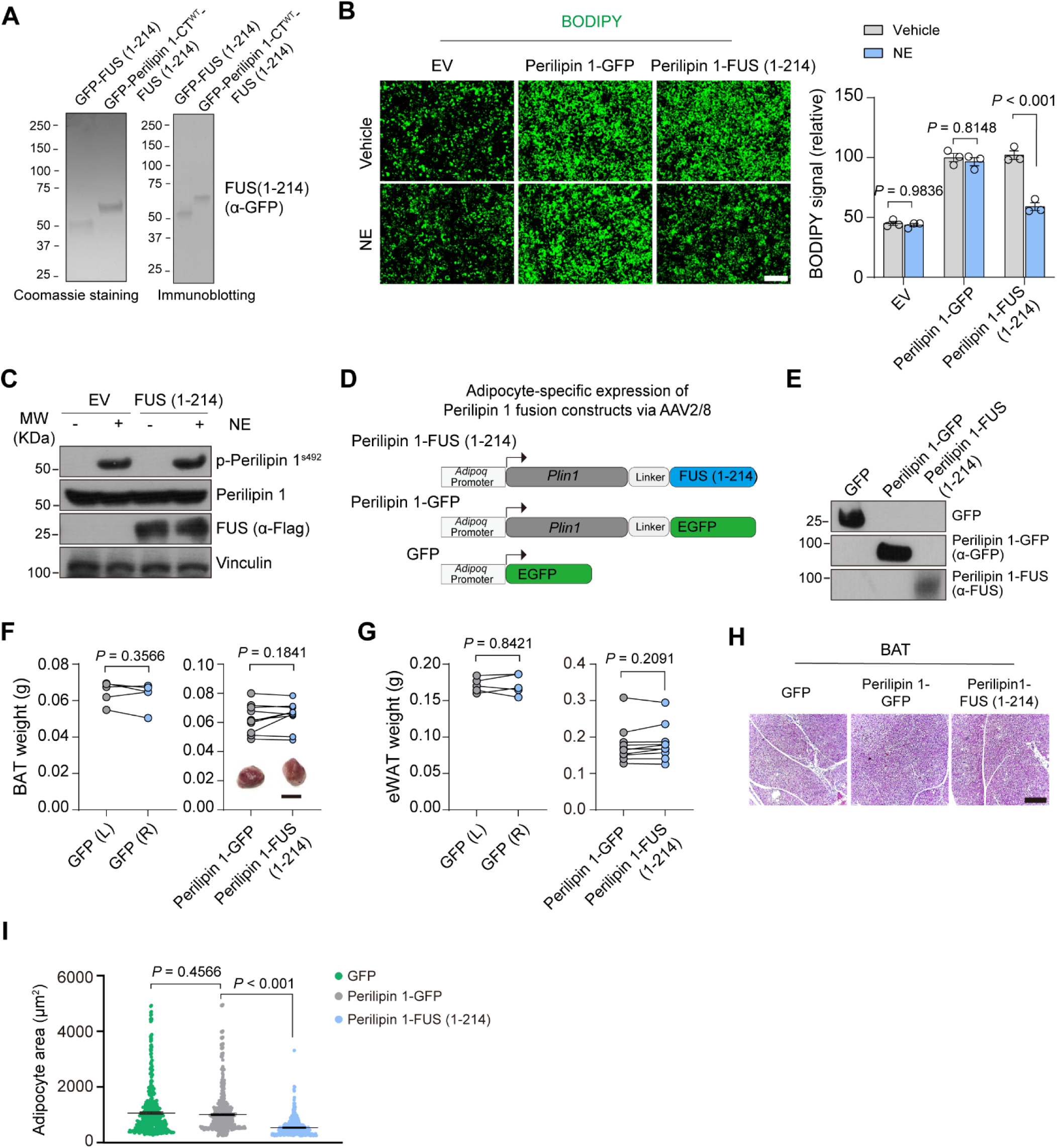
Enhanced perilipin 1 phase separation promotes lipolysis. **A.** <u>Left</u>: Coomassie Brilliant Blue staining of purified GFP–FUS (1–214) and GFP–perilipin 1-CT–FUS (1–214). <u>Right</u>: Immunoblotting with anti-GFP confirmed the protein identity. **B.** <u>Left</u>: Representative BODIPY staining of perilipin 1-knockout adipocytes reconstituted with an empty vector (EV), perilipin 1–GFP, or perilipin 1–FUS (1–214) in response to the treatment with vehicle or norepinephrine (NE) at 0.1 μM for 8 hours. Scale bar: 130 µm. <u>Right</u>: Quantification of BODIPY fluorescence intensity. *n* = 3 per group. Data are presented as mean ± s.e.m. Two-way ANOVA with Šídák’s multiple comparisons test. **C.** Differentiated adipocytes expressing empty vector (EV) or Flag–FUS (1–214) were stimulated with norepinephrine (NE) at 0.1 μM for 10 min. Immunoblot analysis shows phospho-perilipin 1^S492^, total perilipin 1, and FUS. Vinculin was used as a loading control. **D.** Schematic of AAV2/8 vectors driven by the adiponectin (*Adipoq*) promoter to express perilipin 1–FUS (1–214), perilipin 1–GFP, or GFP that were used *in vivo*. **E.** Immunoblot analysis of GFP, perilipin 1-GFP, and perilipin 1–FUS (1–214) in adipose tissue. **F.** Brown adipose tissue weight from matched depots of mice expressing GFP, perilipin 1-GFP, or perilipin 1-FUS (1–214) following bilateral subcutaneous injection. Each data point represents an individual depot, with the left or right depots from the same mouse. Representative BAT images are shown at the bottom inset. *n* = 5 mice for control group, *n* = 10 mice for FUS group. Paired *t*-test. **G.** Epididymal white adipose tissue (eWAT) weight from matched depots of mice expressing GFP, perilipin 1-GFP, or perilipin 1-FUS (1–214) following bilateral subcutaneous injection. Each data point represents an individual depot, with the left or right depots from the same mouse. *n* = 5 mice for control group, *n* = 10 mice for FUS group. Paired *t*-test. **H.** Representative hematoxylin and eosin (H&E) staining of BAT sections from matched depots of mice expressing GFP, perilipin 1-GFP, or perilipin 1-FUS (1–214) following bilateral subcutaneous injection. Scale bar: 210 µm. **I.** Quantification of adipocyte area shown in Fig. 6K. *n* =400 cells per group from 3 mice. One-way ANOVA followed by Tukey’s multiple comparisons test.

**Table S1.**
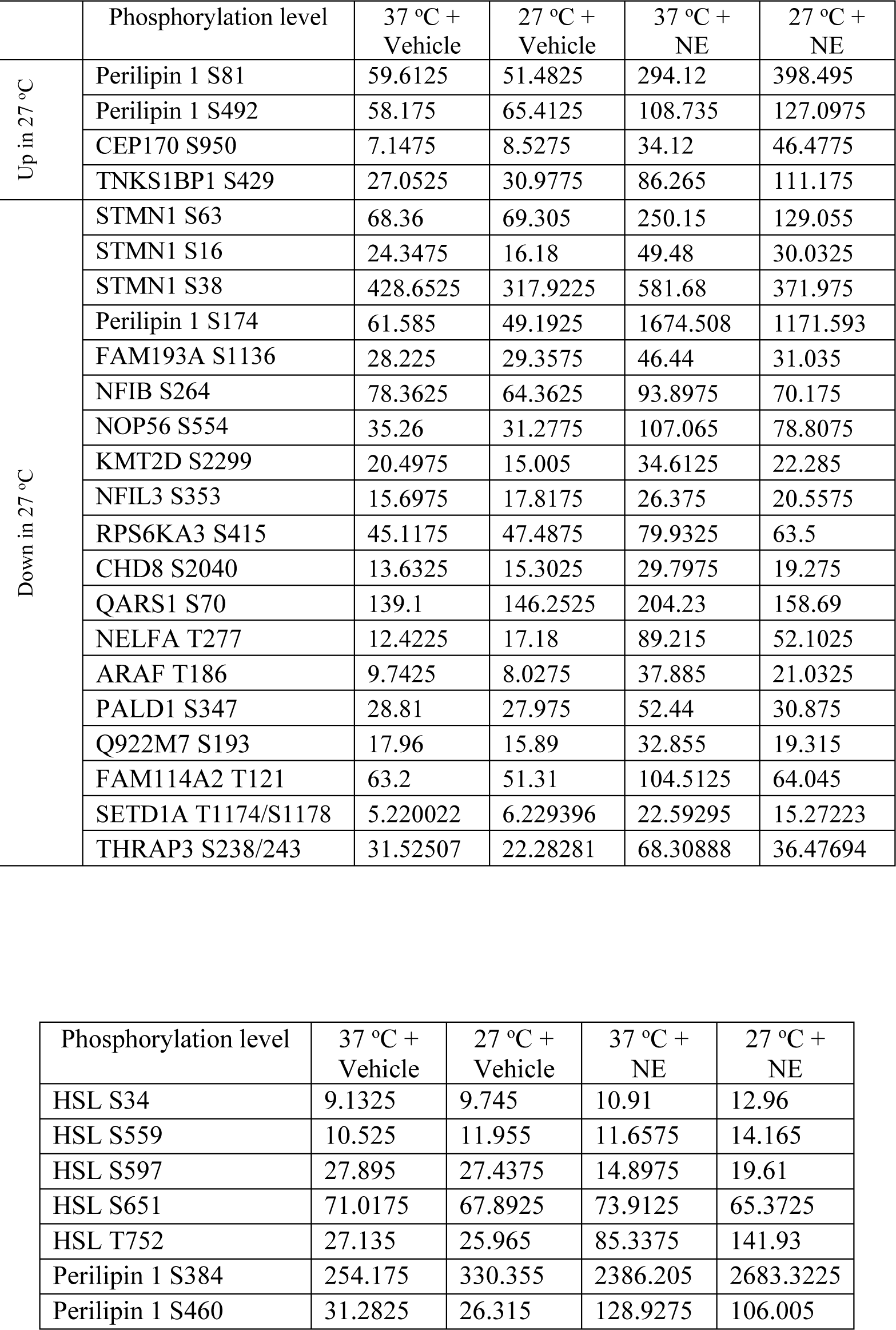
Temperature-dependent regulation of norepinephrine-induced phosphopeptides. Phosphorylation levels of perilipin 1 and HSL

**Table S2.**
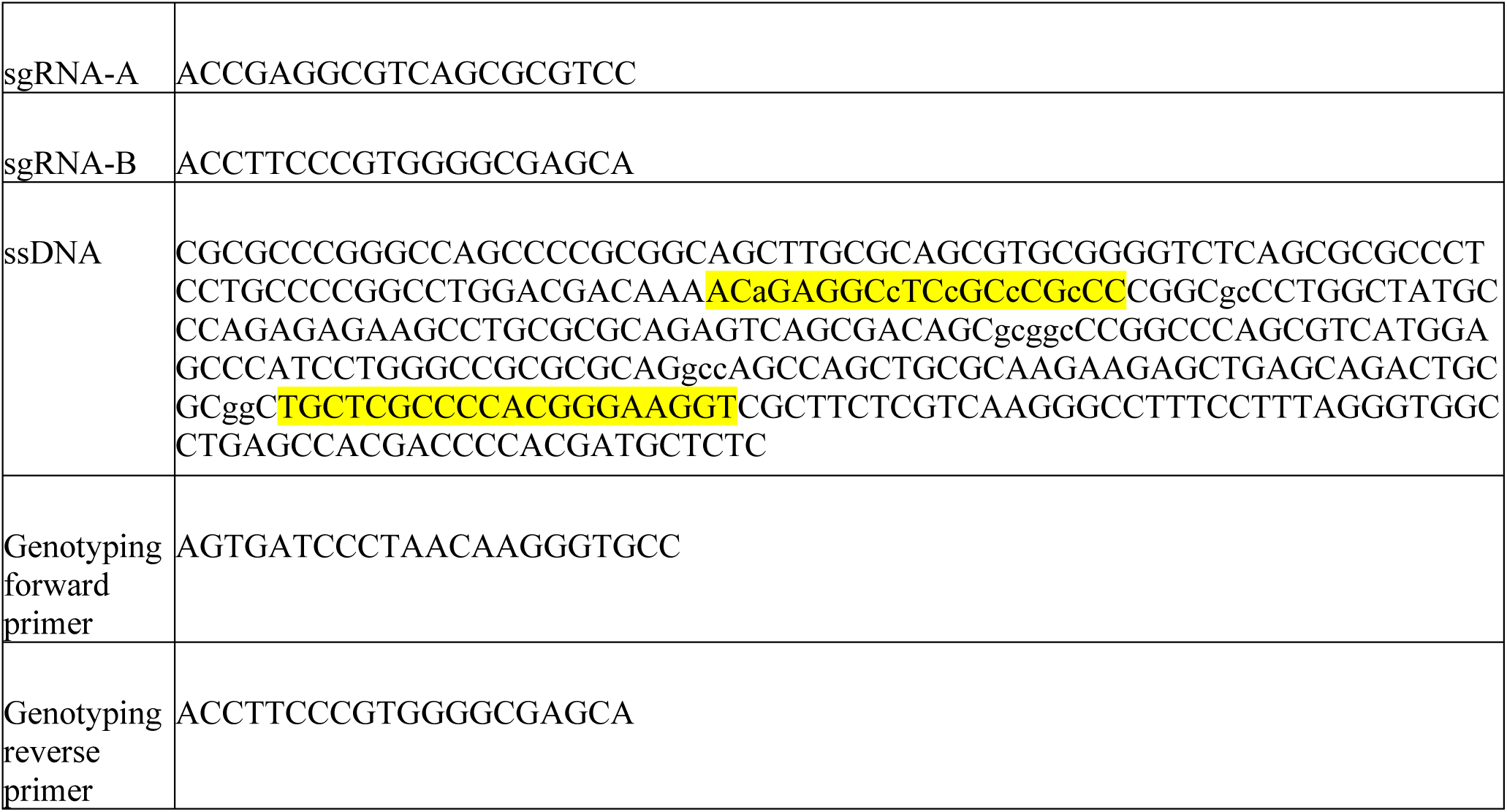
gRNA target sequences and ssDNA donor sequences used for the generation of perilipin 1^FY4A^ knock-in mice.

**Table S3.**
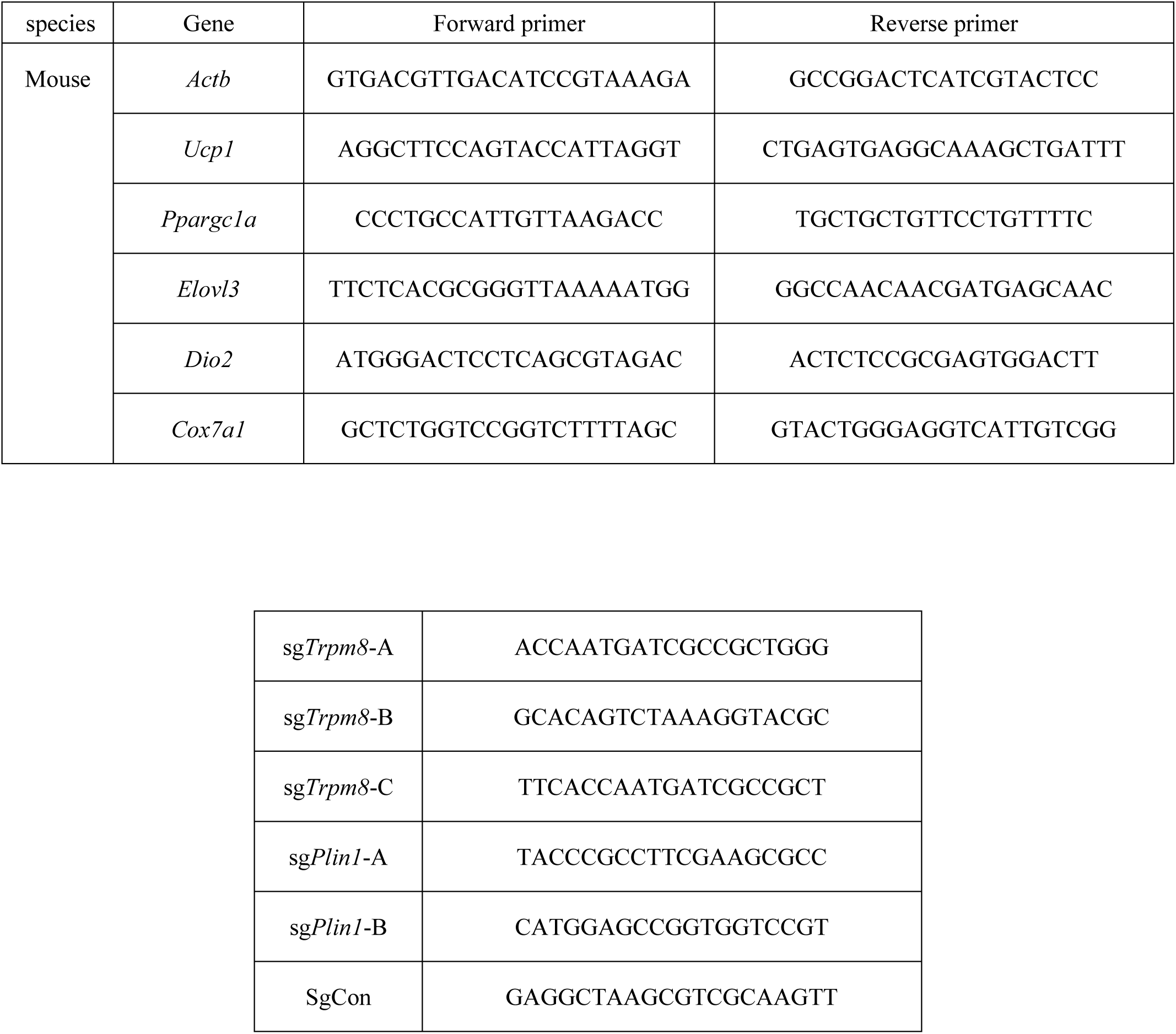
RT-qPCR primer sequences. gRNA target sequences for generation of *Trpm8* and *Plin1* knockout adipocytes

